# Global quantification of mammalian gene expression noise

**DOI:** 10.64898/2026.05.11.724258

**Authors:** Anna Sophie Welter, Florian Mutschler, Mareike Simon, Chiara Giacomelli, Ann-Christin Branscheid, Artür Manukyan, Luiz Gustavo Teixeira Alves, Maximilian Gerwien, Robert Kerridge, Markus Landthaler, Jana Wolf, Matthias Selbach

**Author notes:** these authors contributed equally to the publication (Anna Sophie Welter, Florian Mutschler). Corresponding author: Matthias Selbach, Max Delbrück Center for Molecular Medicine Robert-Rössle-Str. 10, D-13092 Berlin, Germany.

## Abstract

Even cells of the same type growing in the same environment show cell-to-cell differences in protein abundance, a phenomenon known as gene expression noise. This variability can be decomposed into intrinsic components, reflecting molecular randomness, and extrinsic components, arising from differences in cellular state. While gene expression noise has been studied genome-wide in microbes, its global organization remains largely unknown in mammalian cells. Here, we develop a spike-in-based stable isotope single-cell proteomics approach that enables robust quantification of protein-level gene expression noise across thousands of human proteins. We find that protein noise scales inversely with abundance until reaching a plateau, consistent with an extrinsic noise floor and conserved scaling principles observed in bacteria and yeast. Cell cycle stage and cell size contribute substantially to protein variability but do not fully account for the observed heterogeneity. Gene-specific features such as mRNA and protein half-lives and translation efficiency show only weak associations with protein noise, and variability at the mRNA level is a weak predictor of protein variability. Instead, protein noise is largely extrinsic, with coordinated variation across proteins encoding biologically organized cellular states. Consistently, coordinated proteome programs predict intercellular differences in proteome dynamics, linking protein variability to cellular function. Together, these results provide a proteome-wide view of gene expression noise in mammalian cells, establishing that protein-level variability encodes structured and functionally relevant differences in cellular state.

## Main

Cell-to-cell variability in gene expression is a pervasive feature of biological systems, even among cells of the same type in a shared environment ^1,2^. Often referred to as gene expression noise, this variability is not purely stochastic but also reflects differences in cellular state. It can be broadly decomposed into intrinsic components, arising from stochastic molecular processes, and extrinsic components, capturing differences in cellular state, a distinction that provides a useful framework for interpreting gene expression noise ^3^. Gene expression noise is a fundamental property of living systems, enabling phenotypic diversification and adaptive strategies such as bet-hedging ^4,5^. In mammalian systems, noise plays a critical role in diverse biological contexts such as cell-fate decisions during development ^6–10^ and the emergence of drug resistance in cancer ^11–15^.

In bacteria and yeast, genome-wide studies have established that protein noise typically scales inversely with mean abundance until reaching an extrinsic noise floor ^16–18^. In mammalian cells, global measurements of mRNA and protein abundance and turnover at the population level have revealed key principles of gene expression control ^19,20^. However, the genome-wide properties of endogenous protein noise remain unknown, as protein-noise analyses have largely relied on artificial reporter systems that lack proteome-wide coverage ^21–25^. Recently, mass spectrometry-based single-cell proteomics has evolved into a powerful technology to investigate cellular heterogeneity with increasing coverage and throughput ^26–31^. However, inherent challenges such as limited sensitivity, imperfect cross-cell normalization, pervasive missing values, and technical variability have so far hindered robust global quantification of protein-level gene expression noise ^32–35^.

### Single cell spike-in SILAC (scSiS)

We recently showed that spiking in a heavy proteome reference, generated by stable isotope labelling with amino acids in cell culture (SILAC), improves proteome coverage and quantification for low input samples ^36^. We reasoned that this spike-in SILAC (SiS) strategy could offer distinct advantages for single-cell proteomics, including increased proteome coverage, improved comparability across cells, and the ability to directly estimate technical noise. To implement single cell spike-in SILAC (scSiS), we cultivated cells in heavy SILAC medium, prepared a bulk heavy lysate, and spiked it into individual wells of a 384-well plate as an internal reference (Fig. 1A, Extended Data Fig. 1). We then FACS-sorted individual non-labelled (light) cells into each well, processed them, and analysed them on an ultrasensitive mass spectrometer (timsTOF Ultra 2) using a highly sensitive acquisition method (Slice-PASEF) ^37^. Light-to-heavy ratios reflect the abundance of each precursor relative to the shared reference, enabling direct comparison of protein levels across individual cells. Because the heavy reference proteome is constant, these ratios can be placed on a common abundance scale across proteins and cells.

**Fig. 1:**
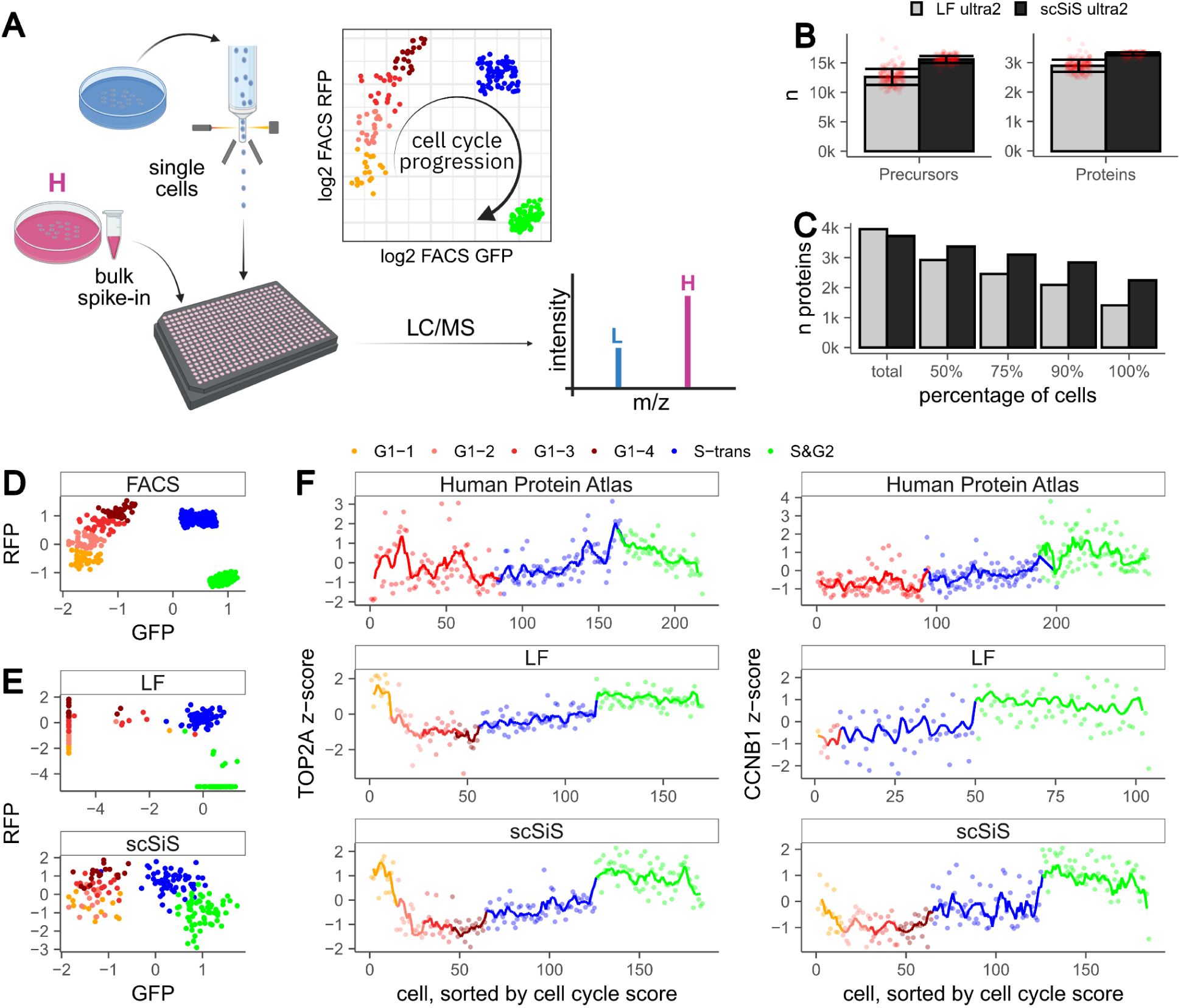
Single cell spike-in SILAC (scSiS) improves proteome coverage and quantification. **A,** Experimental design. U2OS FUCCI cells were grown in light SILAC medium. A separate U2OS population was fully labeled in heavy SILAC medium and used as a common spike-in. The heavy reference was added to 384-well plates, and single light cells were FACS-sorted into each well before digestion and DIA-based mass spectrometric analyses (timsTOF Ultra 2). **B,** Mean number of identified precursors (left) and proteins (right) per run for label-free (LF; 172 cells) and scSiS (185 cells) datasets. Error bars indicate standard deviation; red dots indicate individual runs. **C,** Data completeness, defined as the number of proteins consistently identified across all cells in each sampled subset (total, 50%, 75%, 90% or 100% of cells), for LF and scSiS datasets. **D,** FACS-based z-scores for GFP and RFP fluorescence. **E,** Mass-spectrometry-based z-scored GFP and RFP abundances measured by LF or scSiS. Missing values were set to −5. **F,** TOP2A and CCNB1 profiles across the cell cycle as reported by the Human Protein Atlas (top) and measured by LF (middle) and scSiS (bottom). Points represent individual cells, and lines show a rolling average over six cells.

As a model system, we selected U2OS human osteosarcoma cells expressing a fluorescent ubiquitination-based cell cycle indicator (FUCCI), which has been extensively profiled by single-cell transcriptomics and antibody-based proteomics ^38^. Using a DIA-NN-based ^39^ data processing pipeline, scSiS increased the average number of proteins identified per cell to over 3,000, exceeding the depth achieved by a standard label-free workflow (Fig. 1B). More importantly, scSiS markedly reduced missing values (Fig. 1C). Consistent detection of GFP and RFP enabled scSiS to recapitulate the FACS-based cell cycle assignments more accurately than the label-free data (Fig. 1D–E). Using the GFP and RFP fluorescence we calculated a cell cycle score acting as a pseudo-time (Extended Data Fig. 2A-B). The abundance profiles of several cell cycle-regulated proteins, including topoisomerase II alpha (TOP2A) and cyclin B1 (CCNB1), across the cell cycle score matched those obtained from antibody-based measurements (Fig. 1F). Principal component analysis further revealed clearer separation of cell cycle states in the scSiS data than in the label-free dataset (Extended Data Fig. 2C-D). We conclude that scSiS improves both proteome coverage and quantification of the cell cycle-dependent proteome.

### Global noise scaling

In *E. coli* and yeast, the noise of GFP-tagged fusion proteins scales inversely with protein abundance until reaching an extrinsic noise floor ^16–18^. While mass spectrometry does not directly report protein copy numbers, our protein abundances correlate with protein copy number estimates in U2OS cells (Extended Data Fig. 2E), allowing us to approximate global noise scaling. To assess this relationship, we plotted the log squared coefficient of variation (CV²) against the log mean protein abundance across single cells (Fig. 2A, left). Protein noise decreased with increasing abundance, consistent with intrinsic fluctuations dominating at low expression levels. For highly expressed proteins, noise reached a plateau, indicative of a global extrinsic noise floor (Extended Data Fig. 3A). In *E. coli*, Taniguchi and co-workers estimated global extrinsic noise by correlating expression levels of 13 randomly selected pairs of high-abundance YFP and RFP fusion proteins ^16^. Taking advantage of our proteome-wide measurements, we applied the same strategy to the top 5% (n = 184) most abundant proteins and estimated a global extrinsic noise floor of 16% (95% CI: 14.5 - 17.8%; Fig. 2A left panel, grey lines). This level of variability is substantially lower than estimates reported for *E. coli* (∼30%).

**Fig. 2:**
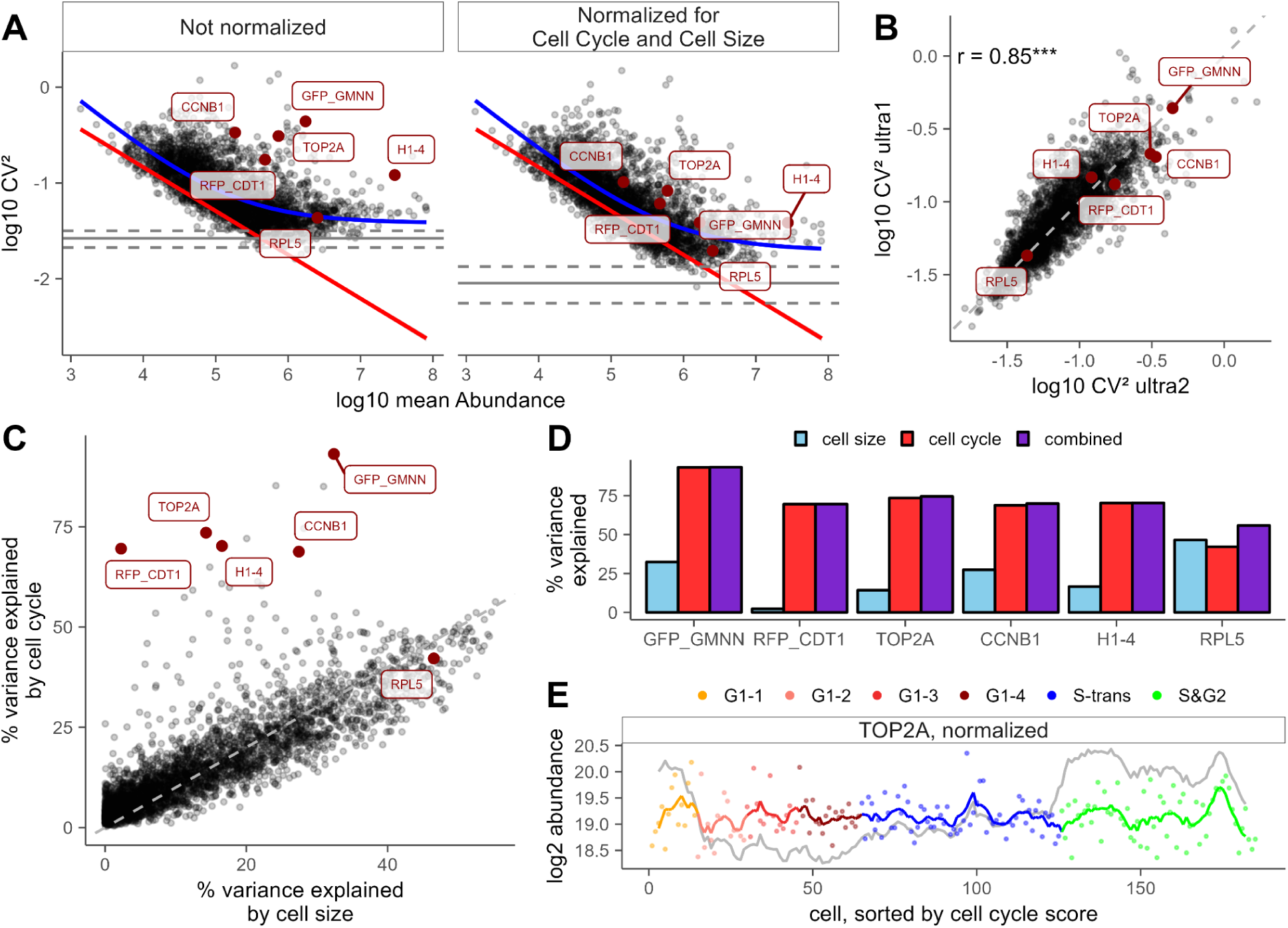
Protein noise follows global scaling with contributions from cell cycle and cell size. **A,** Global noise scaling (CV² versus mean protein abundance). The x axis was scaled to estimated protein copy numbers (Extended Data Figure 2E). Only proteins quantified in more than 100 cells are shown; selected cell-cycle and cell-size-dependent proteins are highlighted. The right panel shows the data after cell-cycle and cell-size normalization (Fig. 2C-E, Extended Data Fig. 3F-H). The blue line shows the best fit to the data (a/μ^β^ + b, Extended Data Fig. 3A), and the red line indicates the linear fit to the technical noise estimate (Extended Data Fig. 3B-E). The solid and dashed grey lines indicate the extrinsic noise floor and its 95% confidence interval, respectively. **B,** CV² of measurements on timsTOF Ultra 1 versus timsTOF Ultra 2. Significance is indicated as follows: * p <= 0.05; ** p < 0.001; *** p < 0.0001. **C,** Variance in the ultra 2 dataset explained by cell cycle or cell size alone (Extended Data Fig. 3F-H). Combining both predictors allows normalization of each protein for cell-cycle- and cell-size-dependent effects. **D,** Variance explained by cell size, cell cycle, or both combined for selected proteins. **E,** Example of TOP2A normalization across the cell cycle. The grey trace shows the original abundance profile, and the coloured trace shows the cell-cycle- and cell-size-normalized profile. The resulting normalized CV² values are shown in the right panel of A.

The observed CV² reflects both biological variation and technical noise. A key advantage of scSiS is that it enables direct estimation of the technical component: because the heavy spike-in is identical across all wells, variability in the heavy channel must arise from technical sources. This allowed us to estimate technical noise for each protein (see Methods). Under conservative assumptions, the estimated technical noise remained consistently lower than the observed total noise across the full abundance range. Importantly, unlike total noise, technical noise showed no visible plateau at high protein levels (Extended Data Fig. 3B–E; linear fit as red line in Fig. 2A), indicating that the observed noise floor arises from biological rather than technical sources. These results indicate that scSiS measurements are not dominated by technical noise and reflect substantial biological variability.

Among the proteins that exhibit a higher noise than expected from the global trend are many canonical cell cycle regulators. This prompted us to ask how much of the observed variability could be explained by cell-cycle stage or cell size. To address this, we used FUCCI-based cell-cycle information and FACS-derived cell size estimates to model protein abundance as a function of these factors (Extended Data Fig. 3F-H). For canonical markers such as cyclin B1, TOP2A, H1-4 and the FUCCI reporters, most of the variance was explained by cell-cycle phase. Other proteins, such as the ribosomal protein RPL5, were better predicted by cell size (Fig. 2C), and combining both predictors further improved model performance (Fig. 2D). Regressing out the contributions of cell cycle and size markedly reduced the variability of cell-cycle-regulated proteins (Fig. 2A, right panel; Fig. 2E, Extended Data Fig. 3G–H) and lowered the estimated extrinsic noise floor to 9% (95% CI: 7.4 - 11.6%). Importantly, many proteins remained more variable than expected even after this normalization, indicating additional sources of noise.

To assess technical reproducibility, we applied scSiS to another set of single cells analysed on a different mass spectrometer (timsTOF Ultra). Cell-cycle traces, global scaling behaviour, and cell-cycle- and size-based effects were consistent with the original dataset (Extended Data Fig. 4). Most importantly, CV² values of individual proteins were highly correlated between instruments, both before and after regressing out cell-cycle and size effects (Fig. 2B; Extended Data Fig. 4J). Together, these findings show that protein noise in mammalian cells follows a scaling relationship shaped by intrinsic fluctuations and a global extrinsic noise floor, extending observations previously reported in bacteria and yeast ^16–18^. Although cell cycle and size contribute substantially to this variability, they do not fully account for the observed protein noise.

### Protein and mRNA noise

Transcriptional bursting is a major source of gene expression noise in mammalian cells ^22,40–42^. However, mRNA levels often fail to predict protein levels, particularly at the single-cell level ^29,38,43,44^. To systematically assess the relationship between mRNA and protein noise, we integrated our data with corresponding single-cell RNA-seq measurements ^38^. We observed a significant, yet modest, correlation between mRNA and protein CV² across genes (Fig. 3A). Since CV² scales inversely with abundance at both the mRNA and protein level, part of the observed correlation arises from the correlation between mRNA and protein abundance itself. To account for this effect, we normalized CV² values (relative CV²) relative to the global mean-CV² relationship (Extended Data Fig. 3A), thereby quantifying noise relative to the abundance-dependent expectation. As expected, this further reduced the correlation between mRNA and protein noise, although the association remained statistically significant (Fig. 3B). The persistence of a significant correlation is notable, given that scRNA-seq and scSiS rely on entirely different technologies, each subject to distinct sources of technical noise.

**Fig. 3:**
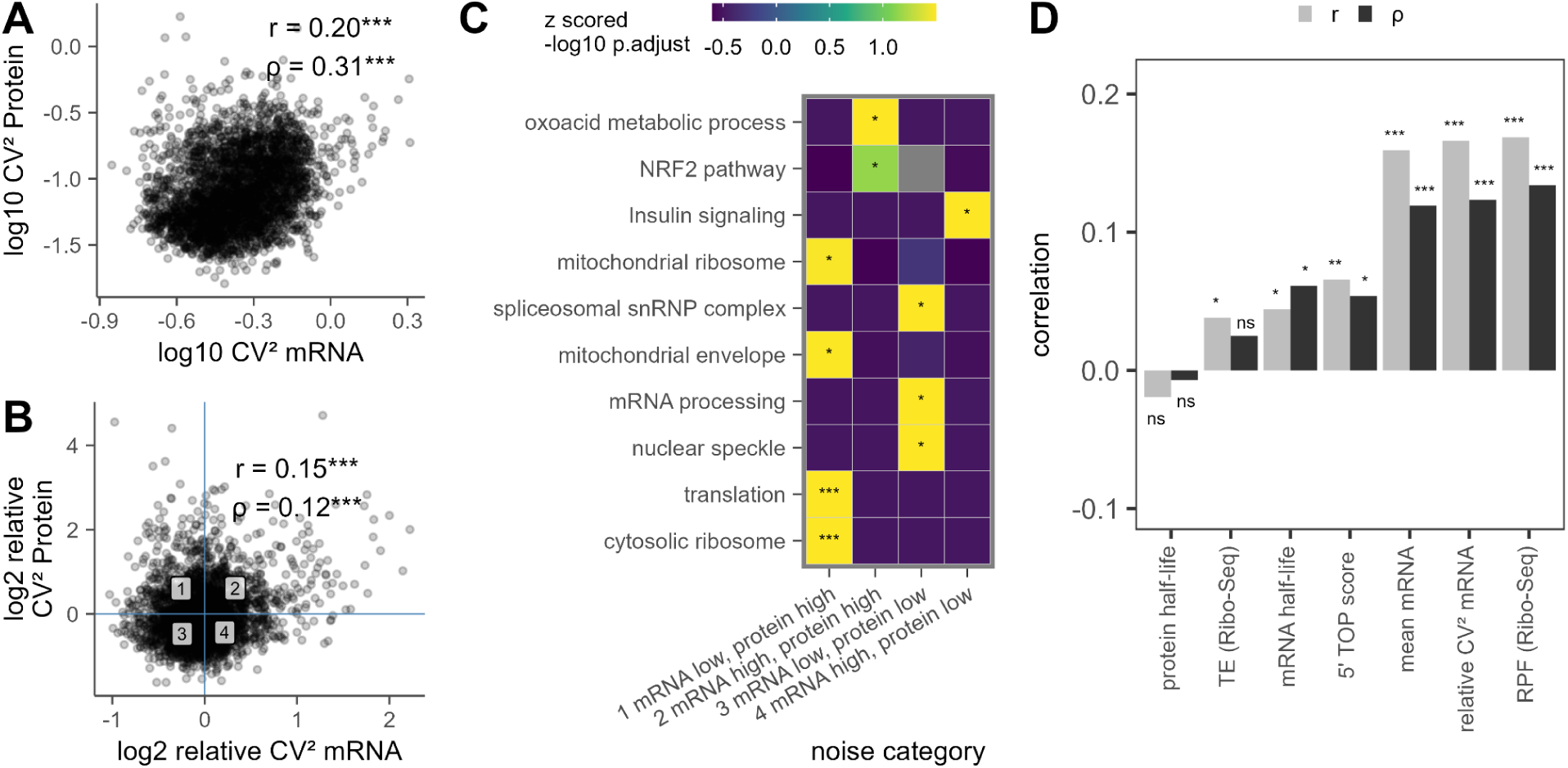
mRNA noise is a poor predictor of protein noise. **A,** CV² of scRNA-seq data versus CV² of single-cell proteomics data. Pearson’s r and Spearman’s ρ are indicated. **B,** Same as in A, but using relative CV² of mRNA and protein. Relative CV² values were calculated as log₂(observed/expected CV²) based on the global mean-CV² relationship (Extended Data Fig. 3A). Blue lines and labels indicate the noise quadrants shown in C. Pearson’s r and Spearman’s ρ are indicated. **C,** Selected enriched GO cellular component (CC), GO biological process (BP), and WikiPathways terms for the noise categories defined in B. Colours indicate z-scored −log₁₀ adjusted p values. **D,** Pearson (r) and Spearman (ρ) correlations between log₂ relative protein CV² and various log₂-transformed features. For all plots, significance is indicated as follows: * p <= 0.05; ** p < 0.001; *** p < 0.0001. For C) BH adjusted p values are shown.

The modest coupling between relative mRNA and protein noise suggests that additional factors shape protein variability beyond transcriptional fluctuations alone. To explore the structure of this variability, we grouped genes according to their combinations of mRNA and protein noise. Strikingly, different noise profiles were enriched for distinct biological compartments, functions and pathways (Fig. 3C). For example, genes with high mRNA and protein noise were enriched for terms related to cellular metabolism, supporting the idea that metabolic heterogeneity contributes to protein noise in this cellular model ^38^. Conversely, genes related to splicing and mRNA processing showed low variability at both the mRNA and protein level, consistent with autoregulatory splicing circuits known to reduce their intercellular heterogeneity ^45^. Also, a couple of mRNA-binding proteins (MBNL1, PUM1, TARDBP, RBM39, ELAVL1) known to autoregulate their own abundance via negative feedback loops ^46–49^ showed reduced protein noise (Extended Data Fig. 5E). Interestingly, mitochondrial proteins and cytosolic ribosomal proteins displayed high protein noise despite relatively stable mRNA levels, suggesting that their variability arises primarily through post-transcriptional mechanisms.

Several factors have been proposed to explain how mRNA fluctuations propagate to the protein level, including mRNA half-life, translation efficiency, translational bursting and protein half-life ^21,23,24,41^. To assess which factors contribute to global protein noise, we performed bulk ribosome profiling (Extended Data Fig. 5A-B), measured protein turnover (Extended Data Fig. 5C), and incorporated previously published mRNA half-life data from U2OS cells ^50^. Additionally, we integrated a 5’ terminal oligopyrimidine (5’ TOP) score as a proxy for mTOR-dependent translation ^51^. We then correlated our relative protein noise with all of these features (Fig. 3D; absolute protein noise in

Extended Data Fig. 5D). mRNA and protein half-lives showed little correlation with protein variability, suggesting that in our model system turnover plays only a minor role on a proteome-wide scale. Total translation output, measured by ribosome footprints, was positively correlated with relative protein noise; however, this relationship largely disappeared after normalizing for mRNA abundance to derive translation efficiency. Although weak, the association between the 5′ TOP score and protein noise is of particular interest. Prior single-mRNA imaging studies have shown that mTOR signaling regulates translational bursting and that 5′ cap-proximal sequences can modulate burst frequency ^23^. Taken together, these results indicate that gene-specific features are weak predictors of protein noise at the proteome-wide level. While this may appear surprising in light of studies using individual reporters, it is consistent with proteome-wide observations in yeast ^17^.

### Noise of abundant proteins is mostly extrinsic

Having found that these features explain only a small fraction of protein noise, we asked whether the observed variability reflects shared extrinsic variation across cells. Extrinsic noise should manifest as coordinated fluctuations affecting multiple proteins simultaneously. To quantify this contribution, we estimated how much of the variance in protein abundances could be explained by shared cellular states captured by the first nine significant principal components (Extended Data Fig. 6A). For highly abundant proteins, these shared components accounted for the majority of the observed variance, indicating that extrinsic noise dominates their variability (Fig. 4A), even after cell cycle and size normalization (Extended Data Fig. 6B). For lower-abundance proteins the fraction explained by shared variation was smaller, likely reflecting a greater contribution of both intrinsic and technical noise. We conclude that extrinsic noise dominates variability for more abundant proteins. This observation extends findings from single-cell transcriptomic studies showing that variability of highly expressed genes largely reflects shared cell-state differences ^52,53^. It also explains why we observed that features associated with intrinsic noise appear to play a minor role (Fig. 3D).

**Fig. 4:**
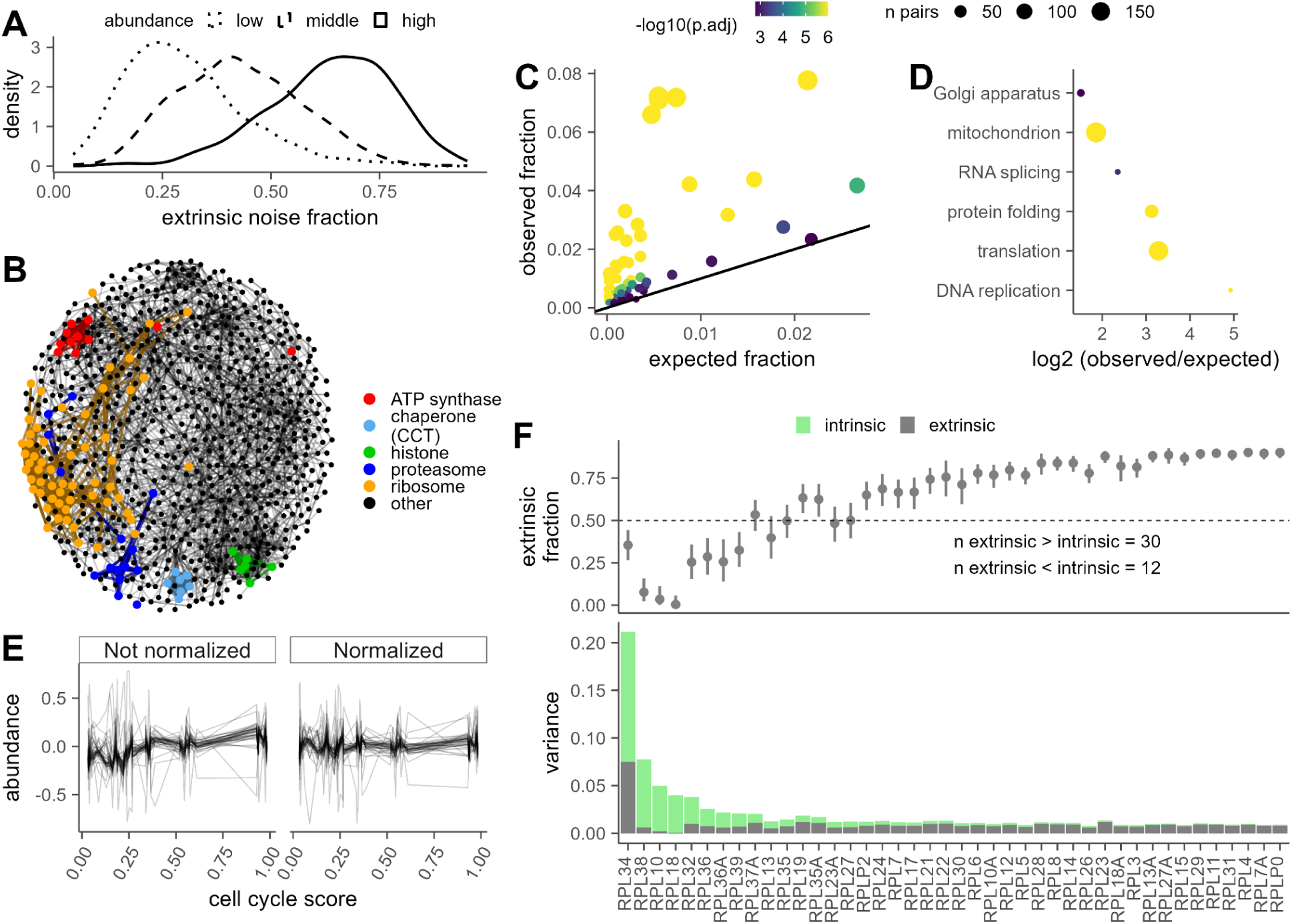
Noise of abundant proteins is largely extrinsic. **A,** Fraction of protein variance explained by shared components (first nine principal components) across proteins, shown for different abundance ranges (lower, middle and upper thirds). **B,** Protein covariance network derived from ridge regression, connecting proteins whose abundance can be predicted from others (R² > 0.5; top 15 predictors with positive coefficients). Selected protein complexes are indicated. **C,** Enrichment of shared Gene Ontology (GO) terms among covarying protein pairs. Each point represents a GO term; the observed fraction of protein pairs sharing the term is plotted against the expected fraction based on the network background. Dot size indicates the number of protein pairs, and colour indicates adjusted p-value. **D,** Selected enriched GO terms, shown as log₂ fold-change between observed and expected fractions. **E,** Median-centered abundance profiles of large ribosomal subunit proteins (RPLs) across cells ordered by cell cycle score, before (left) and after (right) normalization. For better visualization, a randomly sampled subset of cells was used. **F,** Decomposition of total protein noise into extrinsic and intrinsic components for RPL proteins. The upper panel shows the estimated extrinsic fraction (mean ± confidence interval), and the lower panel shows total variance partitioned into extrinsic (grey) and intrinsic (green) contributions.

The large contribution of extrinsic noise implies that proteins involved in related biological processes should fluctuate together across cells. To test this, we extracted covarying proteins in our scSiS data and used them to construct a protein-protein covariance network (see Methods, Extended Data Fig. 6C). Visually, the resulting network recapitulated known cellular modules such as the ribosome, nucleosome, proteasome, chaperonin containing TCP-1 (CCT), and ATP synthase complexes (Fig. 4B), consistent with previous single-cell proteomics studies that used covariance to identify functional protein modules ^26,54^. To test if these patterns reflect genuine biology, we assessed the functional similarity between interacting proteins. Pairs of covarying proteins shared Gene Ontology (GO) annotations far more frequently than expected at random (Fig. 4C). Enriched GO terms included biological processes such as DNA replication, splicing and protein synthesis as well as cellular structures such as mitochondria and the Golgi apparatus (Fig. 4D, Extended Data Fig. 6F). These networks remain enriched for functionally related proteins after normalization for cell cycle and cell size (Extended Data Fig. 6C-F). Remarkably, protein abundances across single cells alone reveal higher-order functional organization of the proteome, even without prior knowledge of pathways or interactions.

One prominent module emerging from the covariance network corresponded to the ribosome. Plotting ribosomal protein abundances across single cells revealed substantial variation, yet most proteins fluctuated in a coordinated manner, consistent with a shared extrinsic driver (Fig. 4E and Extended Data Fig. 6G, left panels). Removing cell-cycle and size effects eliminated the gradual increase in ribosomal abundance across the cell cycle but did not abolish this coordinated variation (Fig. 4E and Extended Data Fig. 6G, right panels). To quantify this relationship we decomposed - for the large and small subunit separately - the noise of each ribosomal protein into an extrinsic (explained variance) and an intrinsic (residual variance) component by using the median abundance of the remaining ribosomal proteins to model the abundance of each individual ribosomal protein. For most ribosomal proteins, the variance explained by this shared signal exceeded the residual variance (Fig. 4F, Extended Data Fig. 6H). These results indicate that ribosomal proteins form a highly coordinated module whose variability is largely extrinsic.

### Extrinsic noise predicts single cell proteome dynamics

The pronounced biological heterogeneity observed in our data prompted us to ask whether differences in proteome states translate into functional differences between individual cells. Notably, ribosomal proteins emerged as a highly variable and coordinated module, suggesting that variability in translational capacity may represent a key axis of cellular heterogeneity. Single-cell proteomics can be combined with dynamic SILAC to quantify protein synthesis at the single-cell level, as recently demonstrated ^31^. To implement this in our spike-in SILAC framework, where the heavy channel is already occupied by the reference proteome, we used medium-heavy SILAC labels to metabolically label newly synthesized proteins for 9.5 h prior to analysis. Using this set-up we quantified between ca. 2,300 and 2,700 proteins in both the preexisting (that is, light, L) and the newly synthesized (that is, medium-heavy, M) form (Extended Data Fig. 7A, B). Reassuringly, the fraction of newly synthesized histone proteins increased progressively during S phase, consistent with their cell-cycle-dependent production during DNA replication (Fig. 5A) and cell cycle reporters showed expected accumulation behaviour during the respective cell cycle phases (Extended Data Fig. 7C).

**Fig. 5:**
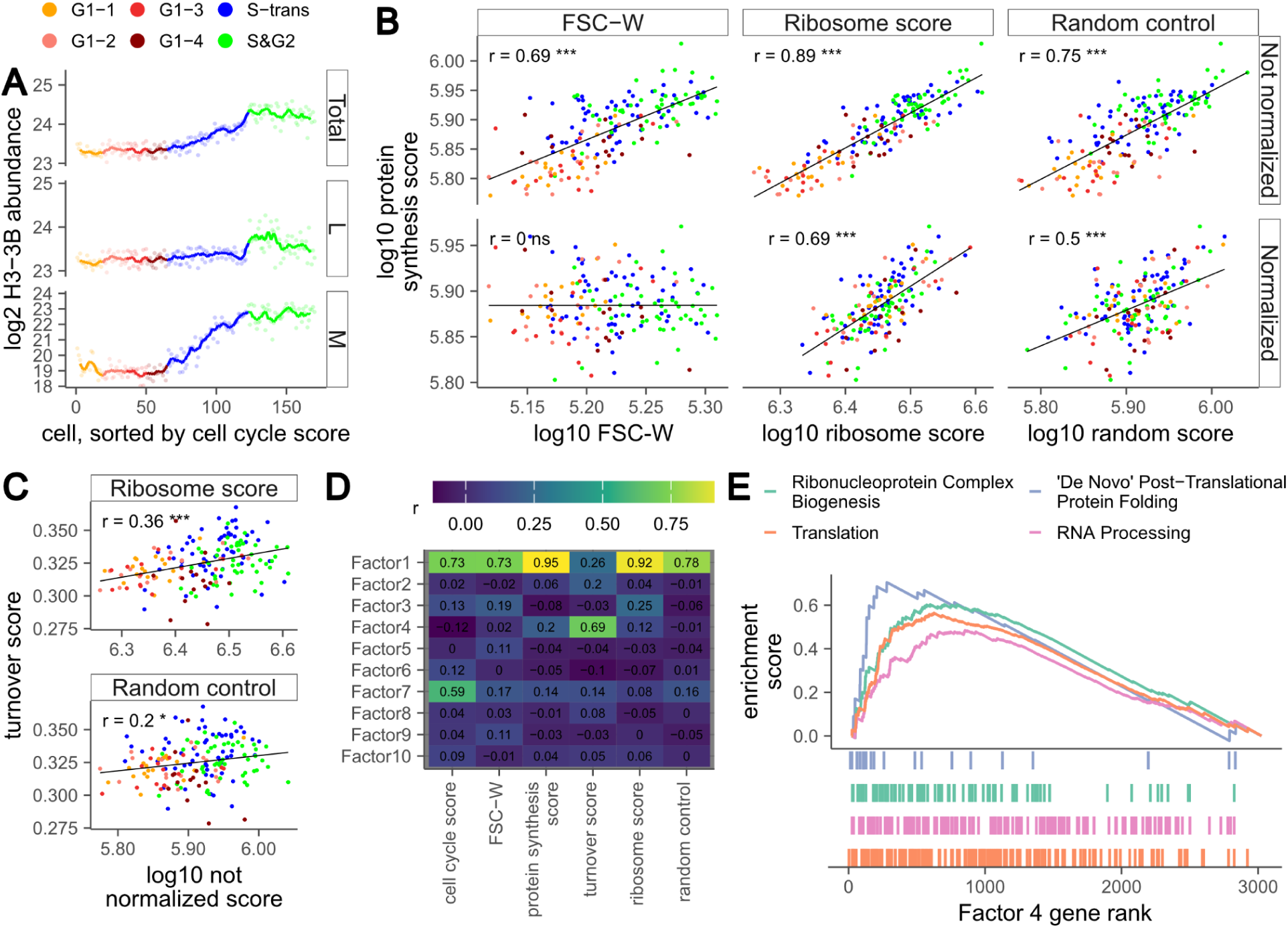
Latent cell states predict proteome dynamics. Cells were pulse-labelled for 9.5 h with medium-heavy SILAC amino acids to distinguish newly synthesized proteins (M) from pre-existing proteins (L), enabling single-cell measurements of protein synthesis and turnover. **A,** Traces of histone H3-3B across the cell cycle, showing log2 abundance of total protein (top), pre-existing protein (L, middle) and newly synthesized protein (M, bottom), coloured by cell-cycle stage. **B,** Protein synthesis score (median abundance of newly synthesized proteins, M) plotted against cell size (log10 FSC-W, left), ribosome score (median abundance of ribosomal proteins, middle), and a random control (randomly sampled non-ribosomal proteins, right) for non-normalized (top) and cell-cycle- and cell-size-normalized data (bottom). **C,** Turnover score (ratio of newly synthesized to total protein abundance, M/(M+L)) plotted against ribosome score (top) and a random control (bottom) for non-normalized data. **D,** Pearson correlation heatmap of MOFA factors with per-cell metadata. Factor signs were aligned so that the strongest absolute correlation is positive. **E,** Gene set enrichment analysis of proteins ranked by their weights on MOFA factor 4, showing selected GO biological process terms. For B and C, significance is indicated as follows: * p <= 0.05; ** p < 0.001; *** p < 0.0001.

To quantify proteome dynamics in single cells, we derived two metrics: a per-cell protein synthesis score and a per-cell turnover score. The protein synthesis score was defined as the median intensity of newly synthesized proteins (medium-heavy channel, “M”) across the 500 most abundant proteins. The turnover score was defined as the sum of all newly synthesized proteins divided by the sum of both newly synthesized and pre-existing proteins (ΣM / Σ(L+M)). For both scores, ribosomal proteins, as well as an equally sized random protein set (serving as a control), were excluded from the data to avoid circularity when relating ribosome abundance to protein dynamics. Cellular protein synthesis increased gradually over the cell cycle and with increasing cell size, but also exhibited substantial cell-to-cell heterogeneity (Fig. 5B, top left panel, Extended Data Fig. 7D, top panel). Regressing out cell cycle and cell size removed this trend, yet substantial heterogeneity remained (Fig. 5B, bottom left panel, Extended Data Fig. 7D, bottom panel). Cellular ribosome abundance (defined as the median abundance of all ribosomal proteins quantified in all cells) was strongly associated with protein synthesis both before and after correction for cell cycle and size (Fig. 5B, middle panels). This observation initially suggests that ribosome abundance predicts translational capacity across single cells. However, similarly strong associations were observed for randomly sampled proteins (Fig. 5B, right panels) or a false protein synthesis score calculated on the pre-existing proteome (Extended Data Fig. 7E), indicating that this relationship largely reflects a global extrinsic state rather than a ribosome-specific effect. In contrast, the turnover score showed weaker correlation with cell cycle, cell size, or randomly sampled proteins, and a moderate association with ribosome abundance (Fig. 5C, Extended Data Fig. 7F). This suggests that turnover captures more specific aspects of proteome regulation that are not dominated by global protein abundance.

To characterize this global extrinsic state more systematically and to identify additional latent sources of variation that may underlie cellular proteome dynamics, we applied Multi-Omics Factor Analysis (MOFA) ^55,56^. MOFA is a latent factor model that decomposes coordinated cell-to-cell variation into a small number of interpretable factors, making it well suited to distinguish broad global effects from more specific proteome programs. Correlation analysis of the MOFA factors showed that Factor 1 was strongly associated with cell cycle, cell size and several global abundance-related metrics, consistent with a broad extrinsic cell state (Fig. 5D, Extended Data Fig. 7G). In contrast, Factor 4 was more selectively associated with the turnover score, suggesting a distinct latent source of variation linked to proteome turnover (Extended Data Fig. 7H). Consistently, gene set enrichment analysis of proteins ranked by their weights on Factor 4 revealed strong enrichment for translation- and ribosome biogenesis-related processes (Fig. 5E). The highest-weighted proteins further support this interpretation (Supplementary Data). For example, DDX21, a central regulator of ribosome biogenesis coordinating rRNA transcription and processing ^57^, showed the strongest contribution to Factor 4. Other prominent contributors included nucleolar proteins such as NOLC1 and NPM1, the translation initiation factor EIF3B, and multiple ribosomal proteins. In addition, proteins involved in proteostasis, such as the ubiquitin-conjugating enzyme UBE2S and the proteasomal ubiquitin receptor ADRM1, were enriched among high-weight proteins, suggesting that this factor captures a coordinated program linking protein synthesis and degradation. Notably, ADRM1 has previously been implicated in regulating cell-to-cell variability in protein turnover ^58^. Collectively, these findings indicate that extrinsic variation in cellular protein abundance is functionally meaningful, capturing biologically organized cellular states and providing predictive insight into cellular phenotypes.

## Discussion

Gene expression noise is a fundamental property of living systems, yet its global organization at the protein level in mammalian cells has remained largely unexplored. Using a stable isotope-based single-cell proteomics approach (scSiS), we provide a proteome-wide view of protein variability in mammalian cells. Unlike reporter- or antibody-based methods that are limited to a small number of proteins, this approach captures thousands of proteins per cell, allowing relationships between proteins to be analysed at scale. Using this framework, we uncover fundamental principles of protein noise in mammalian cells, including conserved scaling behaviour, limited coupling to mRNA variability, and a dominant contribution of shared cellular states.

A central finding of our study is that protein noise in mammalian cells follows conserved scaling relationships, decreasing with abundance until reaching an extrinsic noise floor. Intriguingly, protein variability is only weakly coupled to mRNA noise and poorly explained by gene-specific features such as mRNA and protein half-lives, indicating a partial uncoupling of transcriptomic and proteomic noise. Instead, for more abundant proteins, variability is dominated by extrinsic variation, that is, reflecting different cellular states. Consistent with this, analysis of covarying proteins reveals rich biology, indicative of substantial functional heterogeneity. Indeed, ribosomal and other co-varying proteins define a cellular state that predicts protein turnover of individual cells. Although these relationships are correlative and do not imply causality, they indicate that extrinsic variation in coordinated proteome programs is closely associated with differences in cellular proteome dynamics. Together, these findings support a model in which protein noise reflects structured variation in cellular state rather than purely stochastic fluctuations.

While our approach enables proteome-wide analysis of protein noise, several limitations remain. First, although the use of a shared heavy reference improves quantification, enhances data completeness and enables estimation of technical noise, residual technical variability persists, particularly for low-abundance proteins. As a result, the true coupling between mRNA and protein noise is likely stronger than currently observed. Second, our analysis is restricted to the more abundant fraction of the proteome detectable at the single-cell level. The relative contributions of intrinsic and extrinsic noise for lower-abundance proteins therefore remain unclear, although intrinsic fluctuations are expected to become more prominent at lower copy numbers. Future improvements in proteome coverage will enable these questions to be addressed more comprehensively. Third, because mRNA and protein measurements were obtained from separate cells, their relationship could not be directly assessed at the single-cell level. Future approaches that enable joint quantification of transcriptomes and proteomes in the same cells will be important to resolve how transcriptional variability propagates to protein abundance ^59^. More broadly, the relative contributions of intrinsic and extrinsic noise are likely to depend on cellular context, as suggested by studies in more homogeneous or post-mitotic systems ^60^. Finally, the limited throughput of our method restricts the number of cells that can be analysed and thus the ability to capture rare cellular states. Future developments that combine the quantitative advantages of spike-in strategies with multiplexing approaches may help to overcome this limitation and enable higher-throughput analyses ^61^.

Single-cell proteomics now enables the global quantification of protein-level heterogeneity across thousands of proteins within individual cells, providing a new lens on the structure of gene expression noise in mammalian systems. This opens the door to directly linking proteome-level variability to cellular behaviour, allowing us to explore how differences in proteome states shape responses to perturbations and fate decisions. Extending these approaches to more complex systems, including tissues and *in situ* contexts ^62^, will further reveal how cellular heterogeneity is organized and functions within physiological environments.

## Methods

### Cell culture

U2OS FUCCI cells were kindly provided by the Lundberg laboratory with permission from Hisao Masai. Cells were cultured in DMEM (Gibco; Thermo Fisher Scientific) supplemented with 10% fetal bovine serum (FBS; PAN-Biotech) under standard conditions (37 °C, 5% CO_2_).

For SILAC, cells were maintained in SILAC DMEM (PAN-Biotech) supplemented with 10% dialyzed FBS (PAN-Biotech) that was manually dialyzed an additional time. (ddFBS).

Depending on the label the medium was supplemented with “light”, “medium”, or “heavy” Arginine and Lysine (Arg-0, Arg-6, Arg-10, Lys-0 Sigma Aldrich; Lys-4, Lys-8 Cambridge Isotope Laboratories). Unless otherwise stated, cells were cultured in “light” SILAC medium. For pulse-labelling experiments, cells were switched to “medium-heavy” SILAC medium and incubated for different labelling durations prior to harvest. The durations were 9.5 h for the single cell experiments and 1.5, 4.5, 9.5, and 13.5 h for bulk samples.

For spike-in normalization, a “heavy” SILAC-labelled reference proteome was generated by culturing cells in “heavy” SILAC medium for multiple passages to ensure complete incorporation of heavy labelled amino acids. Labelling efficiency was assessed by mass spectrometry prior to use (Extended Data Fig. 1).

### Cell sorting

Cells were detached using Trypsin-EDTA (Gibco), washed twice with ice-cold PBS, filtered into FACS tubes (Falcon), and kept on ice until sorting. Cell sorting was performed using BD FACSAria III cell sorter directly into 384-well plates (Eppendorf twin.tec) containing 1 µl of lysis buffer (50 mM triethylammonium bicarbonate (TEAB; Sigma-Aldrich), 20% acetonitrile (ACN; CHEMSOLUTE), and 0.1% n-dodecyl-□-D-maltoside (DDM; Thermo Fisher) predisposed into each well.

For experiments including a “heavy” SILAC spike-in, 1 ng of the spike-in standard was included in the lysis buffer.

Gating was performed to exclude debris and doublets, followed by selection of cells at distinct cell cycle stages based on FUCCI reporters.

### Single cell sample preparation

Sorted single cells were subjected to heat at 75 °C for 30 min. Subsequently, 1 µl of digestion buffer containing 2 ng trypsin (Promega) in 50 mM TEAB and 20% ACN was added to each well.

Following digestion, samples were dried and reconstituted in buffer A (3% ACN, 0.1% formic acid (FA; Fluka) in water). Samples were subsequently injected for LC-MS/MS analysis.

### Bulk sample preparation

Bulk sample prep was done for bulk protein turnover, “light” SILAC U2OS for library refinement and the “heavy” SILAC spike-in used to check the labelling efficiency.

Cells were lysed in buffer containing 1% sodium dodecyl sulfate (SDS; Roth) in PBS (Thermo Fisher Scientific) supplemented with protease inhibitors (Roche complete) by boiling at 95 °C for 10 min. Genomic DNA was sheared by sonication using a Bioruptor device (Diagenode) for 10 min.

For protein turnover experiments, proteins were reduced with 5 mM tris(2-carboxyethyl)phosphine (TCEP; Sigma-Aldrich) and alkylated with 20 mM chloroacetamide (CAA; Sigma-Aldrich).

For DDA experiments for copy number estimation proteins were reduced with 10 mM dithiothreitol (DTT; Sigma-Aldrich) and alkylated with 20 mM iodoacetamide (IAA; Sigma-Aldrich). To quench the alkylation the DTT concentration was increased to 20 mM.

Heavy SILAC spike-in samples were not reduced and alkylated to stay in line with the single cell sample preparation.

For all samples, protein clean-up and digestion were performed using SP3 ^63^ with an overnight digest.

### LC-MS/MS analysis

#### Single cell measurements

Single cell measurements as well as 5 ng U2OS bulk measurements for library refinement were carried out on an EASY-nLC 1200 system (Thermo Fisher Scientific) interfaced with timsTOF SCP (library refinement), Ultra (ultra1 data set) and Ultra2 (ultra2 and pulse SILAC data sets) mass spectrometers (Bruker). Chromatographic separation was achieved using an in-house packed column with 75 µm inner diameter and 1.9 µm C18 particles (Dr. Maisch). Mobile phase consisted of buffer A (3% ACN and 0.1% FA in water) and buffer B (90% ACN and 0.1% FA in water). Peptides were resolved over a total runtime of 21 min, including a 14 min gradient segment at a flow rate of 250 nl/min for peptide separation.

The mass spectrometer was operated in data-independent acquisition (DIA) mode using 1-frame Slice-PASEF ^37^. Ion mobility separation happened over TIMS ramps of 100 ms and a range between 0.64-1.45 1/K_0_. A MS1 scan over a range of 100-1700 m/z was followed by 7 MS2 scans ranging from 400-1000 m/z and 15 isolation windows during each TIMS ramp.

#### Bulk measurements

All bulk measurements were done without using the spike-in.

For the bulk data-dependent acquisitions (DDA) used to acquire absolute copy number estimates, samples were measured on an Orbitrap Exploris 480 mass spectrometer. Bulk DIA bulk turnover measurements were acquired on an Orbitrap Astral mass spectrometer. In both cases, the mass spectrometers were coupled to a VanquishNeo system (Thermo Fisher Scientific). The column and mobile phase were identical to those previously described in the single cell measurements.

For bulk DDA, samples were separated over a 33 min gradient with a total run time of 40 min, with a flow rate of 250 nl/min. Full MS scans were acquired in the Orbitrap at a resolution of 60,000 over an m/z range of 350–1600 with a maximum injection time of 10 ms. Data-dependent acquisition was performed using a Top20 method. MS2 spectra were acquired at a resolution of 15,000 with an isolation window of 1.3 m/z and maximum injection time of 22 ms.

Bulk DIA samples were separated over a 23 min active gradient with a 350 nl/min flow rate. MS1 scans were acquired at a resolution of 240,000 over an m/z range of 380-1100 with a maximum injection time of 3 ms. MS2 scans were acquired in 2 m/z isolation windows across a precursor range of 380-980 range with a maximum injection time of 3 ms. Fragment ions were detected between 150-2,000 m/z.

For the labelling check, samples were separated over a 44 min gradient with an EASY-nLC 1200 system (Thermo Fisher Scientific) coupled to a Orbitrap Exploris 480. MS1 scans happened at 60,000 resolution ranging from 350-1600 m/z and in DDA. For the Top20 method an isolation window of 1.3 m/z with a resolution of 15,000 and a maximum injection time of 22 ms was used.

### RiboSeq and RNA-seq

RiboSeq libraries were generated as previously described ^64,65^ with some modifications. Samples were lysed in polysome buffer (20 mM Tris-HCl pH 7.4, 150 mM NaCl, 5 mM MgCl2, 1 mM DTT, 1% Triton X-100, 100 µg/ml cycloheximide and 25 U/ml Turbo DNase (Thermo Fisher Scientific)) and directly frozen in liquid nitrogen. For monosome RNA preparation, total RNA was digested with 2400 U/ml RNase I (Ambion) for 45 min in slow agitation at RT and digestion stopped with 640 U/ml SUPERase•In (Thermo Fisher Scientific). Monosomes were purified by size exclusion with Illustra MicroSpin S-400 HR columns (GE Healthcare) and extracted with 3 volumes of Trizol LS (Thermo Fisher Scientific), chloroform and the RNA Clean & Concentrator kit (Zymo Research). 5 µg of the isolated monosomes were depleted of rRNA using the riboPOOL Ribosome Profiling kit (siTOOLs Biotech). The RNA fragment between 27nt and 30nt size was separated on a 17% Urea PAA gel with markers and their 5′ extremity phosphorylated with 10 U T4 polynucleotide kinase (New England Biolabs) for 1 h at 37 °C. Libraries were generated with the NextFlex small RNA sequencing kit (PerkinElmer), used according to the manufacturer’s instructions. Riboseq library was sequenced single-end on a NovaSeq6000 device (Illumina). Total RNA was extracted from cells lyzed in polysome buffer, as described above. Libraries were constructed using the NEBNext Ultra II Directional RNA Library Prep Kit (New England Biolabs) and sequenced paired-end on a NovaSeq6000 device (Illumina).

#### Raw data processing

Alignment and quantification of Ribo-seq and RNA-seq were performed using custom snakemake (v7.7.0) workflows. Duplicated ribosome protected fragments (RPFs) and FUCCI reads were collapsed and filtered out using a custom perl script. Bowtie (v2.3.2) was used to align deduplicated Ribo and RNA reads to a custom human rRNA reference and remove rRNA reads. All Ribo-seq reads smaller than 26 bps and larger than 30 bps were also filtered out before alignment. For both workflows, Hisat2 (v2.0.0) was used to align remaining Ribo- and RNA-seq reads to a reference of combined human (hg38) and FUCCI marker sequences. RiboTISH (v0.2.5) toolkit was used to generate quality metrics of the uniquely mapped RPFs where we report the fraction of RPF counts in the dominant reading frame out of three frames ^66^. FeatureCounts (v2.0.3) was used to quantify uniquely mapped reads to human gene annotation (GENCODE v39) and FUCCI marker annotations.

### Data Analysis

Generally, data was analyzed with R 4.2.2 and partially R 4.4.2 in R Studio 2025.05.0 and Python 3.13.8 in Jupyter Notebook 4.5.2. The experimental design has been generated with Biorender (Biorender.com) and figures have been polished using Affinity Designer 2 2.3.0. ChatGPT (OpenAI) was used for assistance with statistics, code writing, debugging and language editing. Code and calculations were carefully reviewed and adapted, and all scientific interpretations and conclusions were verified by the authors.

#### Raw data processing

Raw SILAC mass spectrometry files have been processed using DIA-NN ^27,39^ 1.8.2 beta 17. For the analysis of the label-free samples, DIA-NN 2.0 was used. Here, the default DIA-NN settings have been used. For the SILAC data analysis, the following settings have been changed / added: MBR off, –tims-scan, –tims-stack, --fixed-mod SILAC,0.0,KR,label, --lib-fixed-mod SILAC, --channels SILAC,L,KR,0:0;SILAC,H,KR,8.014199:10.008269, --peak-translation, --original-mods, --no-maxlfq. For pulse SILAC samples the medium SILAC masses have been added to the channels command (SILAC,M,KR,4.0251069836:6.0201290267999).

The library used to analyse the true single cell data was generated by refining a spectral library, originally predicted on a human + contaminant fasta file using 5 ng label-free runs measured on timsTOF SCP. The bulk turnover data measured on the Orbitrap Astral was run on a not refined library, based on human and contaminant fasta files.

For the analysis of bulk DDA data, raw files were processed in MaxQuant ^67^ (v2.6.8.0) using default settings with the following modifications: Label-free quantification (LFQ) was performed with a minimum ratio count of 2 and the “Fast LFQ” option disabled. Match-between-runs (MBR) was enabled with a match time window of 0.4 min, match ion mobility window of 0.05, alignment time window of 20 min, and alignment ion mobility window of 1. iBAQ (intensity-based absolute quantification) was calculated with Log fit and charge normalization, Top3 calculation was enabled. The search was performed against the same human proteome and contaminant FASTA as the ones used for the bulk turnover measurements on the Orbitrap Astral.

For the analysis of the DDA labelling check, MaxQuant (v2.6.8.0) was used with the default settings. Raw spectra were checked using Qual Browser Thermo Xcalibur 3.1.66.10 (Thermo Fisher Scientific).

#### Processing of DIA-NN output

Processing of the scSiS files was done as described in the pipeline published in Welter *et al* (2024) ^36^. Briefly, all scSiS files were filtered using `diaSiS::clean_DIANN` as well as additionally for Contaminants in the Protein.Ids column. SiS data was further processed using `diaSiS::filter_DIANN` with CalCols = “any” to use Precursor.Translated or Ms1.Translated if only one of them is available. For pulse SILAC experiments, numberChannels was set to 3. Precursor ratios were calculated using `diaSiS::calculate_SILAC_ratios` with useFilter =“standard”, precursorPerProtein = 1, globalReference = “H”, globalCalculation = “all” and PGcalculation = “sc”. Additionally, we filtered for precursor ratios of log10(L/H) (no pulse and pulse data) and log10(M/H) (only pulse data) < 0.5 to further remove contaminants before calculating protein ratios and abundances. Finally, runs were filtered for a median log10 L/H ratio per < 0.

Label-free data was filtered for Precursor.Charge > 1, Lib.PG.Q.Value < 0.01, Lib.Q.Value < 0.03 and Contaminants in Protein.Ids. Here protein intensities were calculated based on the mean log10 Precursor.Normalised instead of relying on the PG.MaxLFQ column of the DIA-NN output.

Additionally to the described steps, all true single cell data (label-free and scSiS) was manually filtered for IDs (> 3000 SILAC, ultra2; > 2700 SILAC, ultra1; > 2700 label-free, ultra2; and > 2300 L & M in pulse), and based on PCAs of 100% complete proteins. Additionally, summed intensities per run were calculated and outliers visually removed.

The bulk protein turnover measurements on the Thermo Astral were analyzed using the same DIA-NN version as the single cell data. They were post-processed using a different in-house pipeline described in Dueren *et al.* (2026) ^68^ that does not require the presence of a spike-in.

#### Rescaling of scSiS protein intensities

To scale scSiS abundances to protein copy number estimates we used intensity-based absolute quantification (iBAQ) ^19^ from bulk DDA measurements on an Orbitrap Exploris 480 system (see above).

The proteinGroups.txt file from the MaxQuant analysis was filtered to remove potential contaminants and reverse database hits. To ensure data robustness, the median iBAQ value across five biological replicates was calculated for each protein group. Absolute protein copy numbers per cell were then estimated using the proteomics ruler approach ^69^. This method leverages the stoichiometric constancy between genomic DNA mass and histone proteins to provide an internal standard for scaling MS signals. For these calculations, we assumed a hyperdiploid genome for U2OS cells (62 chromosomes ^70^, ≅ 8.6 gigabase pairs) and a nucleosome footprint of 197 bp, yielding a theoretical expectation of 8.73*10^7^ copies per core histone type. A scaling factor was derived by dividing this theoretical value by the summed median iBAQ values of all detected H2A isoforms; this factor was then applied to all iBAQ values to arrive at cellular copy number estimates.

Finally, we aligned these bulk estimates with the scSiS heavy-reference scale. To this end, we computed the median histone H/L ratio across G1 cells, which was about 4.35, reflecting the excess of the heavy spike-in relative to a single cell. To account for this, we multiplied the bulk copy number estimates by this factor. We then fitted the relationship between the rescaled copy-number estimates and the measured heavy-reference intensities using a zero-intercept linear model (Extended Data Fig. 2E). This model was used to predict the heavy-reference intensity for each protein, and scSiS protein abundances were obtained by multiplying the measured L/H ratio by the predicted reference intensity.

#### Rescaling of scRNAseq Data

We reanalysed the published Smart-seq2 U2OS FUCCI dataset (GSE146773) from Mahdessian *et al*. (2021) to obtain ERCC-normalized RNA abundances while preserving cell-to-cell differences in total cellular mRNA. Raw endogenous and ERCC spike-in counts were filtered as following: removal of cells with very low library size or few detected genes and exclusion of wells with aberrant ERCC behaviour (zero ERCC totals, unusually low ERCC totals or ERCC / endogenous ratios, or excessively high ERCC fraction). ERCC-based size factors were computed with `scuttle::computeSpikeFactors` and applied to obtain ERCC-normalized counts, which correct for capture / amplification efficiency but retain biological variation in total RNA per cell.

#### Cell Cycle Score

For each sample, raw GFP and RFP fluorescence readouts from FACS were log2-transformed and used to fit a two-dimensional ellipse capturing the joint distribution of signals (Extended Data Fig. 2A). The center of this ellipse was used as a reference to calculate per-cell polar coordinates, where angles were derived from normalized GFP and RFP offsets. To linearize the distribution, we identified the largest gap in the angle distribution and rotated all angles such that the gap aligned with zero. The transformed angles were then scaled to a continuous score ranging from 0 (early G1 phase) to 1 (late G2 phase).

#### Cell Cycle Models

To normalize for cell cycle and cell size, we utilized the cell cycle score and the FACS forward-scatter (FSC-W). Both were log-transformed and scaled. While cell-size changes were assumed to have a linear effect on protein abundance, the cell cycle score was encoded in linear, squared, cubic, sine and cosine terms to capture different forms of protein behaviour over the cell cycle. For each protein group, we fitted linear models describing protein abundance as a function of (i) cell size (FSC-W), (ii) cell cycle features alone, or (iii) cell size and cell cycle features combined. The potential overfitting of model ii) and especially iii) was intentionally exploited in order to be conservative regarding the remaining variance explained by other, unknown features. To assess the significance of each model, we generated a null distribution by randomly permuting protein abundances within each protein group and repeating the modeling procedure.

#### Modelling of Mean-Noise Relationship

The relationship between log CV² and log mean protein abundance (Fig. 2A, Extended Data Fig. 3A) was fitted assuming three different models: i) an inverse dependence without limit but allowing a variable scaling exponent (CV² = a/μ^β^), ii) an inverse dependence with a lower limit but without a variable scaling exponent (CV² = a/μ + b), and iii) a generalized model with inverse dependence with limit and variable scaling exponent (CV² = a/μ^β^ + b). Based on AIC and BIC, iii) outperformed both other models and was therefore used to calculate the relative noise measure (relative CV²) for all data sets (colour gradient in Extended Data Fig. 3A).

#### Noise Floor Estimation

Global noise floors (grey lines in Fig. 2A and Extended Data Fig. 4I) were estimated using pairwise co-fluctuations between the top 5% high abundant proteins, adapting the method described in Taniguchi *et al.* (2010) ^16^. Briefly, protein abundances were assumed to be modulated by a shared multiplicative factor across runs. Under this model, the normalized co-fluctuation between two proteins x and y,

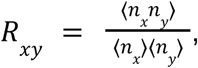

is expected to equal 1+CV². Hence, the noise floor was estimated as *R*_*xy*_ – 1, averaged across all protein pairs. We converted intensities to linear space prior to analysis and computed pairwise *R*_*xy*_ values over all runs in which both proteins were detected. The global floor was defined as the mean across all protein pairs. Confidence intervals were obtained by bootstrap resampling of runs.

#### Technical noise

Technical noise was estimated under the assumption that all variability in the H channel is purely technical, whereas variability in the L channel consists of technical and biological variance. We also assumed that technical noise is predominantly intensity-dependent and behaves independent between channels. Since mass spectrometry errors in practice are not independent and not all variability in the H channel is caused by mass spectrometry but can also happen upstream, these assumptions lead to a more conservative estimate.

To model technical variability, precursor-level L and H intensities were first normalized across runs using median-based normalization relative to a reference run, based on shared precursors between runs. The reference run was selected so it had the highest number of precursors. The resulting normalized intensities were log10-transformed, and only precursors observed in at least 70 measurements were retained. Mean log10 H intensities (*H̅*) were calculated per precursor and split into equally sized bins. The mean variance of log10 H intensities (var(*H̅*)) was computed within each bin to capture the intensity dependence of technical noise. An exponential function *Var*(*H̅*) = a · *e^−b·H̅^* + *c* was fitted using non-linear regression (nls) to describe the relationship between log-intensity and log-variance across the intensity bins (Extended Data Fig. 3B). This model was then used to predict intensity-dependent technical variance for the L channel. Assuming independence of technical variance between L and H, the total technical variance of precursor L/H ratios was estimated as the sum of the observed H variance and the predicted L variance. Using this technical error estimate, we simulated measurements for 500 runs. Therefore, we calculated the mean log10 L/H for each precursor and calculated a normal distribution around that mean with the individual technical variance. 500 ratios per precursor were randomly drawn from that distribution. Since the precursor coverage per protein group is sparse, we also included peptide-specific dropout rates based on the real data (Extended Data Fig. 3C). That way, a precursor only detected in 80% of experimental runs would only be detected in 80% of simulated runs. The simulated ratios were then aggregated to protein abundances using the same pipeline as for our experimental data. Since the DIA-SiS pipeline uses two different intensity types (Ms1.Translated and Precursor.Translated) to aggregate to protein level, normalization and modelling were done for each intensity type individually.

For pulse-labelled data, the estimation was done for the L and M channels separately.

#### Turnover

The calculation of protein turnover and half-lifes followed the approach established in Schwanhäusser *et al.* (2011) ^19^ and Baum *et al.* (2019) ^71^. For each protein group, log H/L (nascent / old proteins) values were aggregated across replicates per timepoint by taking the median. Linear regression without intercept was used to calculate protein turnover based on the slope. Protein degradation constants (kdp) were then derived by correcting these slopes for cell division based dilution effects. To estimate cell cycle duration of our cells, we followed the approach of Leduc *et al.* (2025) which assumed that core histones are predominantly diluted by cell division instead of degradation ^72^. Specifically, protein turnover rates of canonical histones were used to calculate the cell division time as *T*_*cc*_= *log*(2)/*turnover*_*histone*_. The cell cycle duration was obtained by averaging across histone-derived estimates, leading to a mean T_cc_ of 31.64 h (95% CI: 25.83 < 31.64 < 37.47). Protein half-lifes (hf) were calculated as *hf* = *log*(2)/*kdp*.

#### Extrinsic Noise Modelling

##### Global noise decomposition

To globally decompose total protein variability into intrinsic and extrinsic components, we used a PCA-based approach, assuming that the first components will reflect major cellular states (e.g. PC1 reflecting cell size and cell cycle; Extended Data Fig. 2C-D, Extended Data Fig. 3G-H) causing shared variation across cells. Parallel Analysis was used to determine the significant numbers of components per data set (9 for not normalized, 13 for normalized). We then performed a linear regression between the abundance of each protein and these components. The variance explained by this model was interpreted as the extrinsic component while the remaining variance was interpreted as the intrinsic noise component.

##### Complex-specific noise decomposition

Noise decomposition was performed separately for the small (RPSs) and large (RPLs) ribosomal subunits on the non-normalized data. For each protein, a complex score was defined as the median abundance of all ribosomal proteins within the same subunit, excluding the protein of interest. A linear model was then used to predict abundance of each individual protein from the corresponding complex score. The variance explained by the fitted values was interpreted as the extrinsic variability component, whereas the residual variance was taken as the intrinsic variability component.

#### Network analysis

Protein-protein relationships were inferred using ridge regression. For each protein, its abundance across samples was modeled as a function of the abundance of all other proteins across the cells using regression as implemented in the `glmnet` package. Models were fitted using 100-fold cross-validation to determine the optimal regularization parameter (λ), α was set to 0 in order to run as Ridge regression.

Model performance was assessed using cross-validated R², and only proteins with R² > 0.5 (not normalized data) or R² > 0.4 (normalized data) were kept for downstream analysis. For each target protein, the top N predictors (only passing the R² threshold; N = 15) were selected based on the value of their regression coefficients. Only positive regression coefficients were taken into account.

A protein interaction network was constructed in which nodes represent proteins and edges represent predictive relationships inferred from the regression models. Edges were drawn from predictor to target protein.

As quality control and to identify functional modules within the network, a GO enrichment (hypergeometric test on GO terms) was performed. To avoid bias, any connection between two proteins was considered to be bidirectional. For each protein, GO terms were pulled and the number of observed proteins-pairs sharing a GO term were calculated. For the background, the number of theoretical possible protein-pairs sharing a GO term was computed combinatorically. P-values were Benjamini-Hochberg adjusted.

#### Ribosome, Turnover, and Translation Score

Per-cell ribosome scores were calculated using the subset of ribosomal proteins detected in all cells. For each cell, the score was defined as the median total abundance, where the total abundance was computed as the sum of L and M abundances.

A per-cell random control score was computed analogously by randomly sampling the same number of proteins used for the ribosome score from the subset of proteins with complete coverage across all cells, with ribosomal proteins excluded from the sampling pool.

Per-cell translation scores were calculated using proteins detected in all cells, excluding those used for the ribosome score and the random control. From this pool, the 500 most abundant proteins - ranked by their median total abundance across cells - were selected. The translation score for each cell was then defined as the median M abundance of these proteins.

To compute per-cell turnover scores, proteins used for the ribosome score and random control were excluded. The remaining M and total protein abundances were transformed to linear space. For each cell, M and total abundances were summed separately, and the ratio of summed M to summed total abundance was calculated, yielding the medium-heavy fraction of the observed proteome.

## Data and code availability

All scripts to reproduce figures and process data as well as all necessary processed data is available via Zenodo (DOI: 10.5281/zenodo.20067990). The mass spectrometry proteomics data have been deposited to the ProteomeXchange Consortium via the PRIDE partner repository with the dataset identifier PXD078157. RiboSeq data will be made available via GEO.

## Acknowledgements

We thank Hisao Masai (Tokyo Metropolitan Institute of Medical Science) for permission to use the U2OS-FUCCI cell line, and Emma Lundberg (KTH Royal Institute of Technology) for providing the cells. We thank Hans-Peter Rahn and Kirstin Rautenberg (MDC FACS facility) for assistance with FACS sorting, Jeannine Wilde and Madlen Sohn (BIH/MDC Genomics Technology Platform) for sequencing support, and Christian Sommer for support with mass spectrometry. We are grateful to Fabian Coscia, Di Qin, Melissa Klingeberg (all MDC) and Christoph Krisp (Bruker) for fruitful discussions on single-cell proteomics, and to Vadim Demichev (Charité) and Henrik Zauber (MDC) for helpful discussions and suggestions on single-cell proteomics data analysis.

This work was supported by the German Ministry of Research, Technology and Space (BMFTR) via the national research node for mass spectrometry in systems medicine MSTARS (16LW0240) and the ERANET Neuron consortium ALTRUISM (01EW2001) to MSe. LGTA and ML were supported by the Helmholtz Association Initiative and Networking Fund through the project “Virological and immunological determinants of COVID-19 pathogenesis – lessons to get prepared for future pandemics” (KA1-Co-02, COVIPA).

## Author Contributions

ASW: Formal analysis, Investigation, Methodology, Software, Visualization, Data Collection, Data Curation, Writing - Original Draft Preparation

FM: Formal analysis, Investigation, Methodology, Data Collection, Sample Preparation, Visualization, Writing - Original Draft Preparation

MSi: Investigation, Consultation, Writing - Original Draft Preparation

CG: Investigation, Sample Preparation

ACB: Data Collection, Sample Preparation

AM: Data Collection

LGTA: Data Collection

MG: Data Collection, Sample Preparation

RK: Sample Preparation, Software

ML: Funding acquisition, Supervision

JW: Funding acquisition, Consultation, Supervision

MSe: Conceptualization, Investigation, Consultation, Funding acquisition, Project administration, Supervision, Writing - Original Draft Preparation

FM & ASW contributed equally and have the right to list their name first in their CVs.

