## Extended Data Figures for "Global quantification of mammalian gene expression noise"

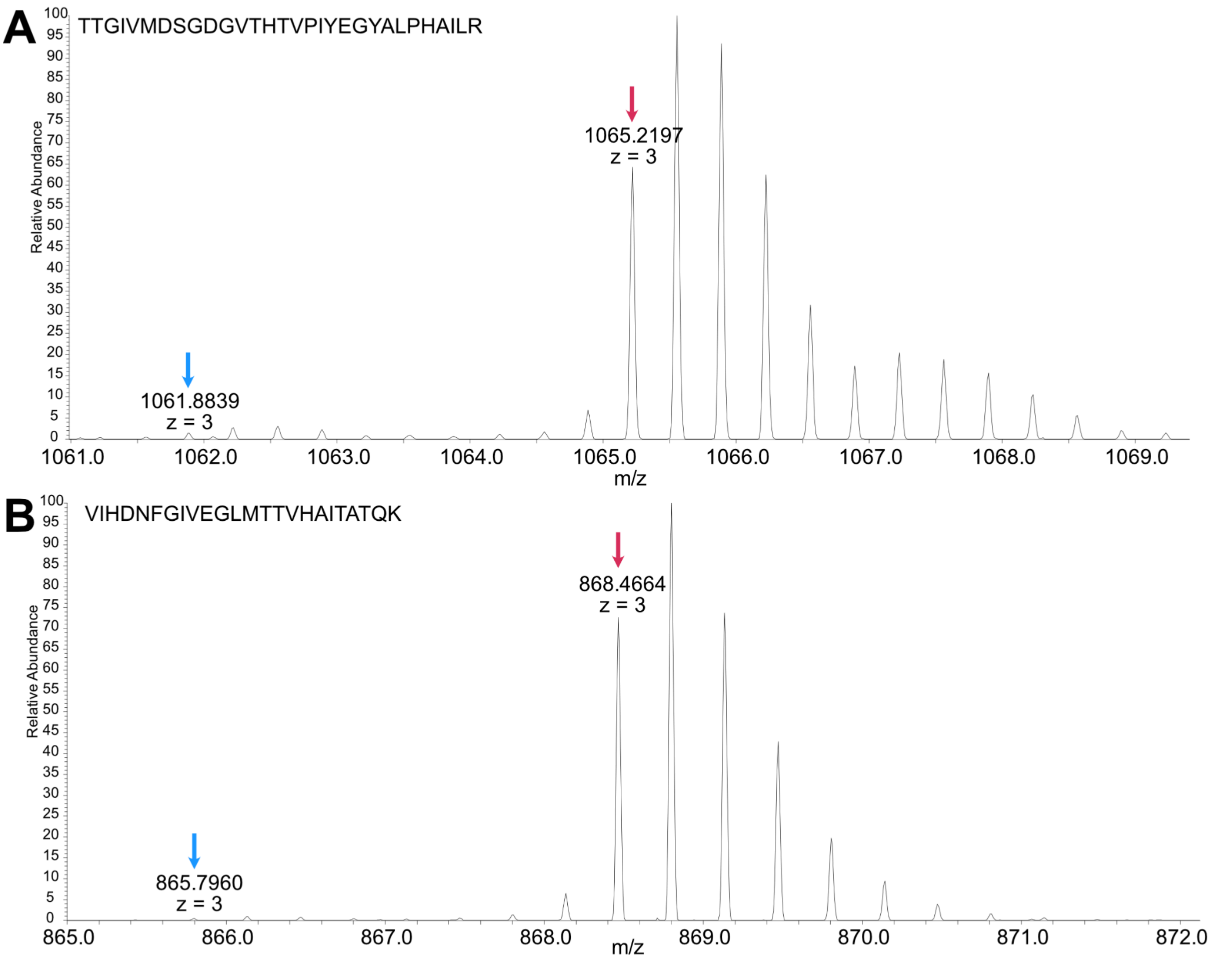


**Extended Data Fig. 1 | Labelling efficiency of the heavy SILAC reference.**

U2OS FUCCI cells were cultured in medium containing heavy arginine ($^{13}C_6{}^{15}N_4$-Arg; Arg10) and lysine ($^{13}C_6{}^{15}N_2$-Lys; Lys8) for several generations to generate a fully labelled internal reference for spike-in SILAC. Representative MS1 spectra of the heavy reference show an arginine **(A)** and a lysine **(B)** containing peptide. Red and blue arrows indicate the m/z of the monoisotopic peaks of the heavy and light peptides, respectively.

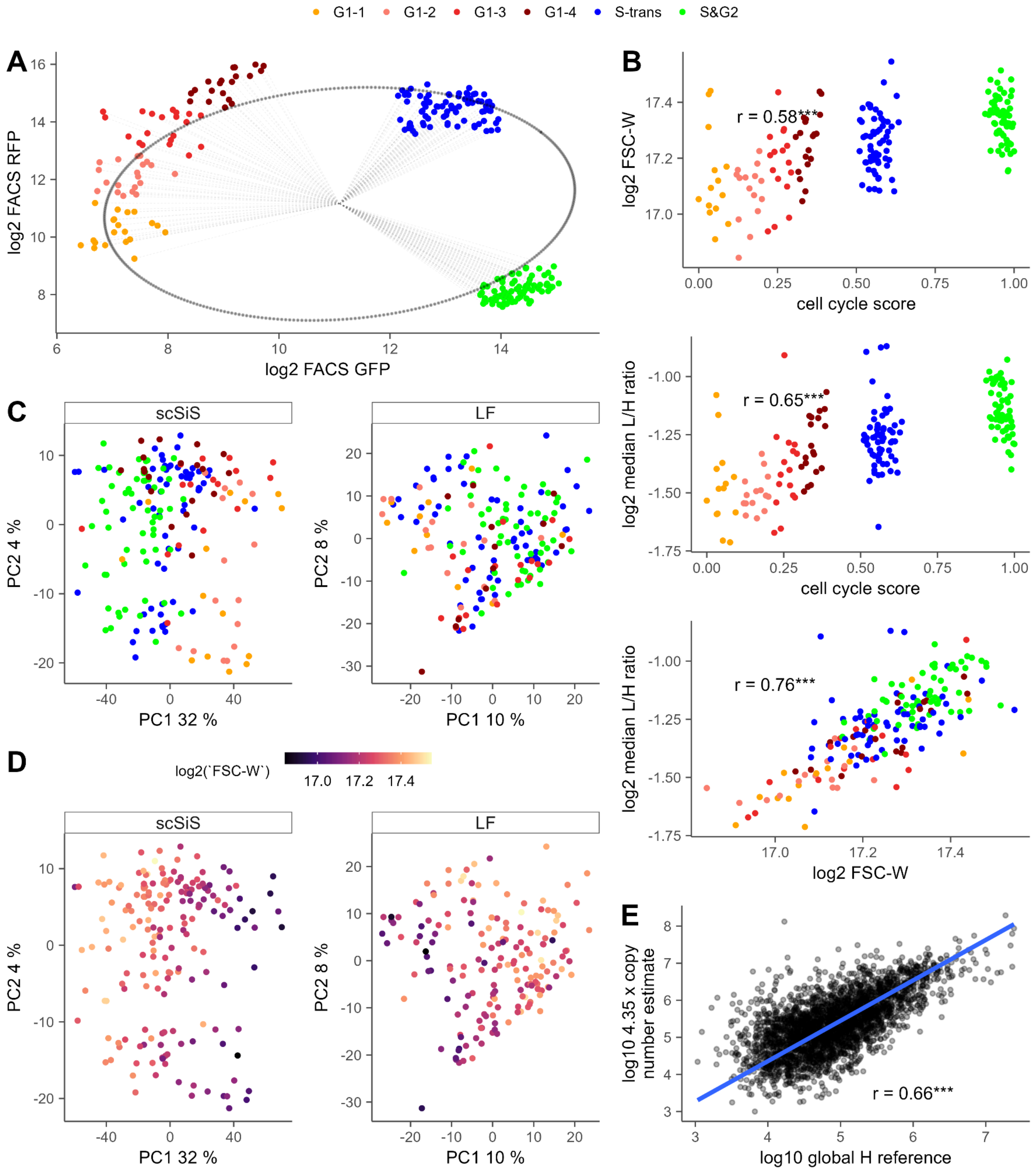


**Extended Data Fig. 2 | Cell state inference and abundance scaling.**

A | Determination of a cell cycle score from FUCCI fluorescence.

Scatter plot of GFP and RFP fluorescence measured by FACS for U2OS cells expressing the FUCCI cell cycle reporter used for single-cell proteomics. Each point represents one cell, colored by cell cycle stage (see legend). An ellipse was fitted to the distribution, and the angular position of each cell relative to the ellipse center (grey dashed lines) was used to derive a continuous cell cycle score for downstream analyses.

B | Relationship between cell cycle score, cell size, and total protein abundance.

Top panel, correlation between the inferred cell cycle score and the FACS forward scatter width (FSC-W), which serves as a proxy for cell size. Middle panel, correlation between the cell cycle score and the median $\log_2$ light/heavy (L/H) ratio across proteins for each cell, reflecting total cellular protein abundance measured by scSiS. Bottom panel, correlation between median $\log_2$ L/H ratio and $\log_2$ FSC-W. Together, these measurements show that both FSC-W and the median L/H ratio capture variation in cell size across the cell cycle.

C–D | Principal component analysis of scSiS and label-free proteomics data.
PCA of proteins quantified without missing values in the scSiS and label-free (LF) datasets. Points represent single cells. Cells are colored by C) inferred cell cycle stage, and D) FACS forward scatter width (FSC-W), a proxy for cell size.

E | Calibration of the heavy reference abundance scale.
Correlation between the log10 global heavy reference abundance and log10 protein copy number estimates times 4.35 (due to the median L/H ratio of core histones in G1 cells) derived from bulk U2OS proteome measurements using intensity-based absolute quantification (iBAQ). The bulk data were acquired by data-dependent acquisition (DDA) on an Orbitrap Exploris 480 mass spectrometer. The fitted relationship (blue line, forced through 0) was used to rescale the heavy reference and to derive protein abundance estimates for the single-cell data.

For all plots, significance is indicated as follows: *: $p \le 0.05$; ** $p < 0.001$; *** $p < 0.0001$.

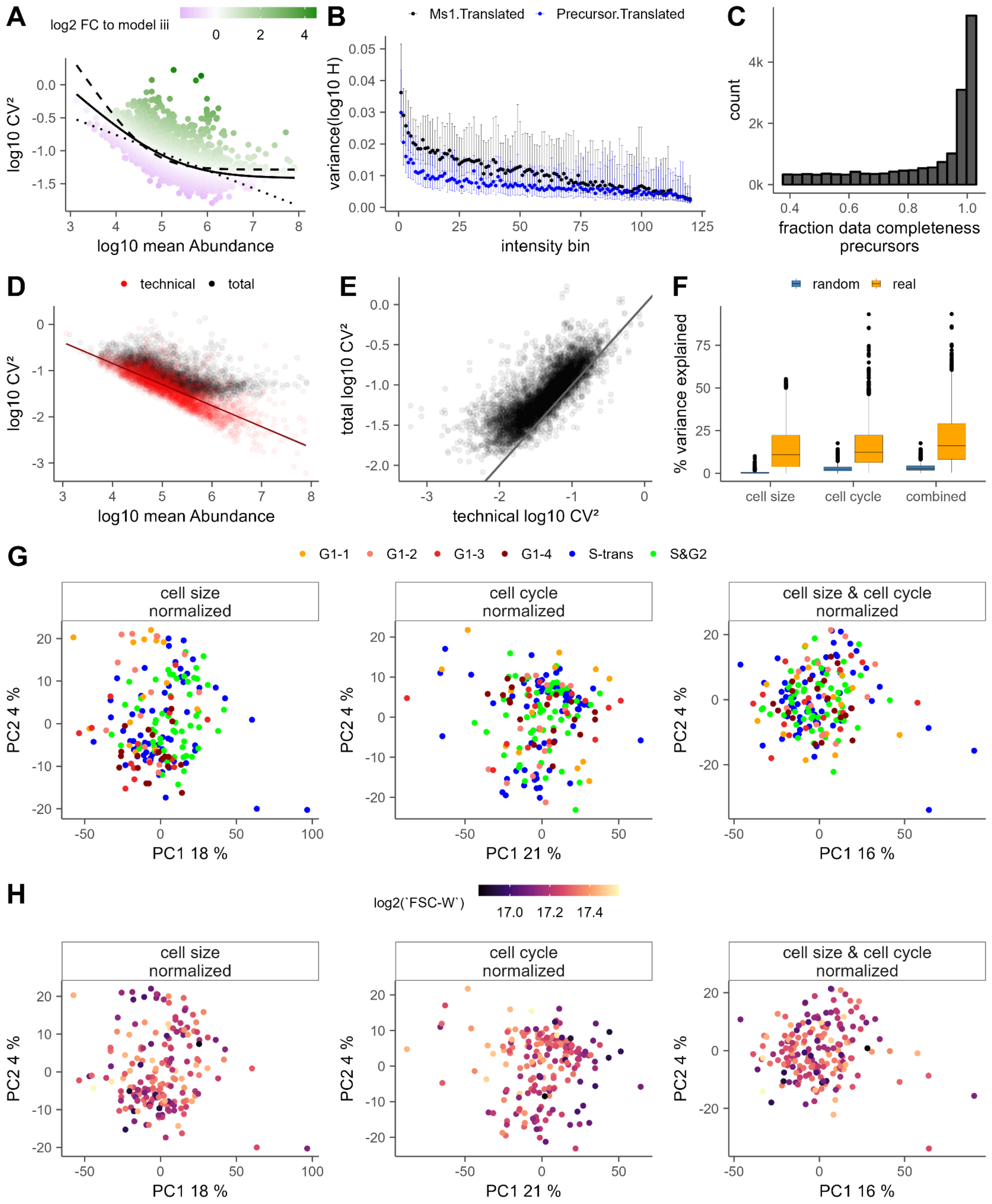


**Extended Data Fig. 3 | Modelling of total CV$^2$, technical CV$^2$ and cell cycle and cell size effects of timsTOF Ultra 2 measurements.**

A |Modelling of the mean–noise relationship and calculation of ΔCV².

The relationship between log10 CV² and log10 mean protein abundance was fitted using three models: (i) a power-law model with variable scaling exponent ($CV^2 = a/\mu^\beta$; dotted line), (ii) a model with an inverse dependence and constant noise floor ($CV^2 = a/\mu + b$; dashed line), and (iii) a generalized model allowing a variable scaling exponent ($CV^2 = a/\mu^\beta + b$; solid line). Models incorporating a noise floor (ii and iii) provided a better fit than the power-law model based on AIC and BIC. The generalized model (iii), which relaxes the assumption of a fixed inverse scaling, was the best performing one and therefore used to calculate relative $CV^2$ as the log2 fold-change relative to the expected noise at a given abundance (colour scale).

B-E | Estimation of technical noise.

To estimate technical variability, the log-variance of heavy precursor intensities was calculated for both MS1-level signals (Ms1.Translated, black) and precursor-level signals (Precursors.Translated, blue) and grouped into 120 equally sized intensity bins. For each bin, the mean log-variance (dots; error bars indicate quantiles) was used to model intensity-dependent technical noise (see Methods). This model was then used to simulate light-channel variability, which was combined with the observed heavy-channel variance to estimate total technical variance for each precursor. Based on this, synthetic L/H ratios were generated and processed identically to the real data to obtain protein-level intensities. Data completeness was taken into account when aggregating precursor signals to proteins (panel C). The resulting technical $CV^2$ values were consistently lower than the observed total $CV^2$ (panels D, E). The linear fit through the technical noise (panel D) corresponds to the red line shown in Fig. 2.

F | Performance of cell cycle and cell size regression models.

FACS-derived fluorescence measurements and forward scatter were used to regress cell cycle and cell size effects from protein abundances across single cells, independently for each protein. Randomized models were generated as controls. While random models explained little variance, the real models captured a substantial fraction of variability, with a median explained variance of ~10–16% depending on the model.

G–H | Principal component analysis after regression of cell cycle and cell size effects.

Protein abundances were normalized using residuals from models correcting for cell cycle, cell size, or both. Principal component analysis of the normalized data shows a loss of structure associated with cell cycle (G) and cell size (H), with little to no remaining grouping by these factors (see Extended Data Fig. 2 for non-normalized data).

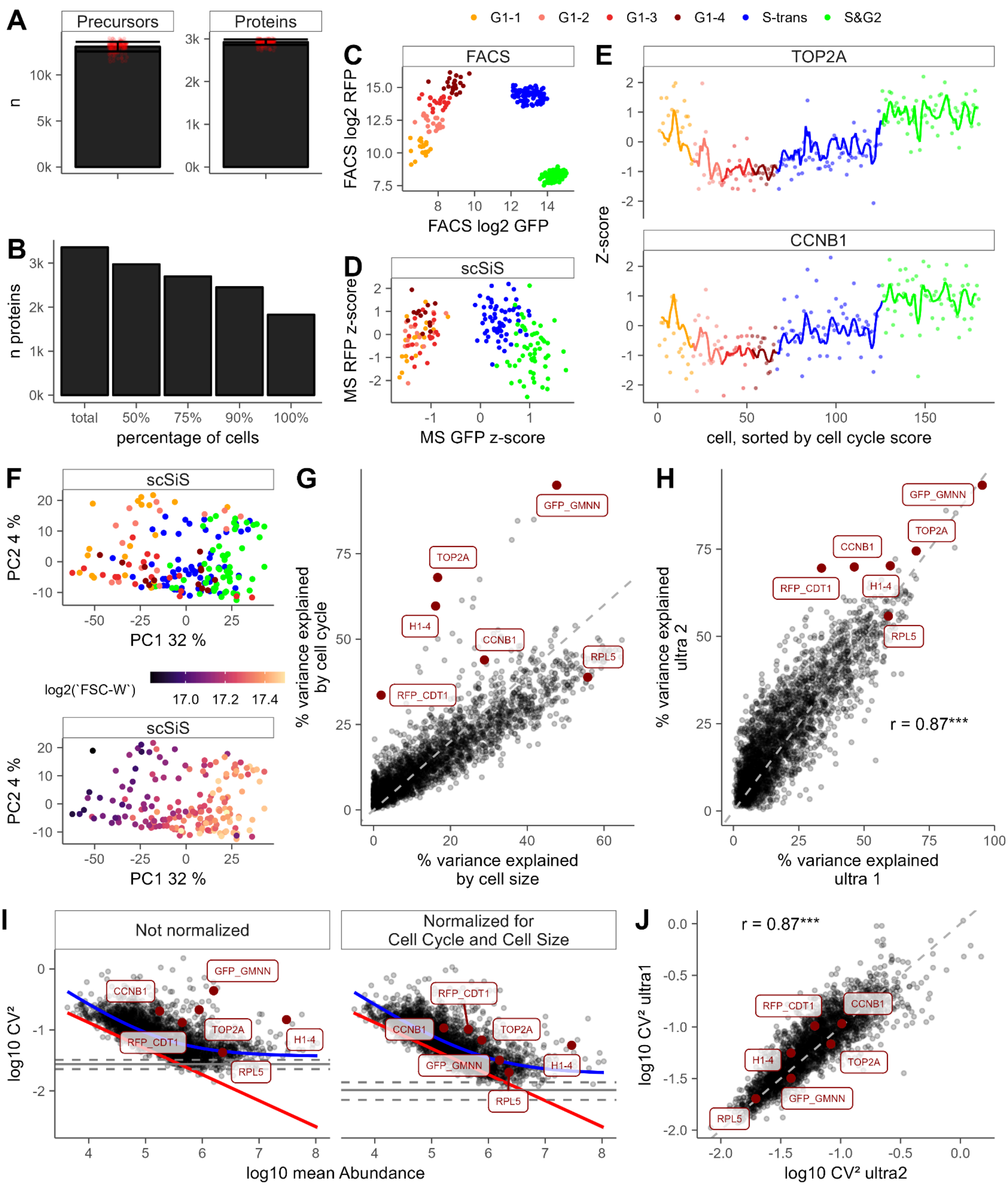


**Extended Data Fig. 4 | Quality and CV$^2$ of timsTOF Ultra 1 measurements.**

<u>A-B | Number of identifications and data completeness</u>.
Mean number of precursors and proteins (panel A) and their data completeness (panel B) across all 180 single cell measurements. Error bars indicate standard deviation, red dots are individual runs. Y axis in thousands.

<u>C-D | FACS fluorescence and MS based protein quantities of cell cycle markers.</u>

FACS measured fluorescence of GFP and RFP (panel C) and z-scored GMMN-GFP and CDT1-RFP intensities measured with the ultra 1. Missing values have been set to -5. Colours indicate cell cycle state.

E | Traces of TOP2A and CCNB1 across the cell cycle according to mass spectrometry.
The points indicate individual cells, lines indicate the rolling average over 6 data points. Cells were sorted by cell cycle score.

F | Principal component analysis of the ultra 1 data set.
PCA of proteins quantified without missing values. Points represent single cells. Cells are colored by inferred cell cycle stage (top), and FACS forward scatter width (FSC-W), a proxy for cell size (bottom).

G | Cell cycle and cell size normalisation of the ultra 1 data set.
Variance of each protein in the ultra 1 data set that can be explained by cell cycle or cell cycle alone. Each dot represents one protein. A few selected cell cycle and cell size dependent proteins are highlighted.

H | Explained variance for of the ultra 1 and the ultra 2 data sets.
Comparison of the variance explainable for every protein by combining cell size and cell cycle information for the ultra 1 and ultra 2 data scSiS data sets with a pearson correlation of 0.87.

I | Global noise scaling properties of the ultra 1 data set.
X axis has been rescaled (Extended Data Fig. 2E) to reflect copy number estimates better. Only proteins with data completeness > 100 cells are shown. Right plot before cell cycle and cell size normalisation, left plot after cell cycle and cell size normalisation. The blue line reflects the best fit through the data ($a/\mu^\beta + b$, Extended Data Fig. 3A) and the red line indicates average technical noise. Solid and dashed greys line indicate the extrinsic noise floor (15-18 % not normalised and 8-12 % normalised) and the 95% CI, respectively.

J | Comparison of the $CV^2$ of cell cycle and cell size normalised data.
$CV^2$ (cell cycle and cell size normalised) of measurements on timsTOF ultra 1 versus timsTOF ultra 2 with a pearson correlation of 0.87.

For all plots, significance is indicated as follows: *: $p \le 0.05$; ** $p < 0.001$; *** $p < 0.0001$.

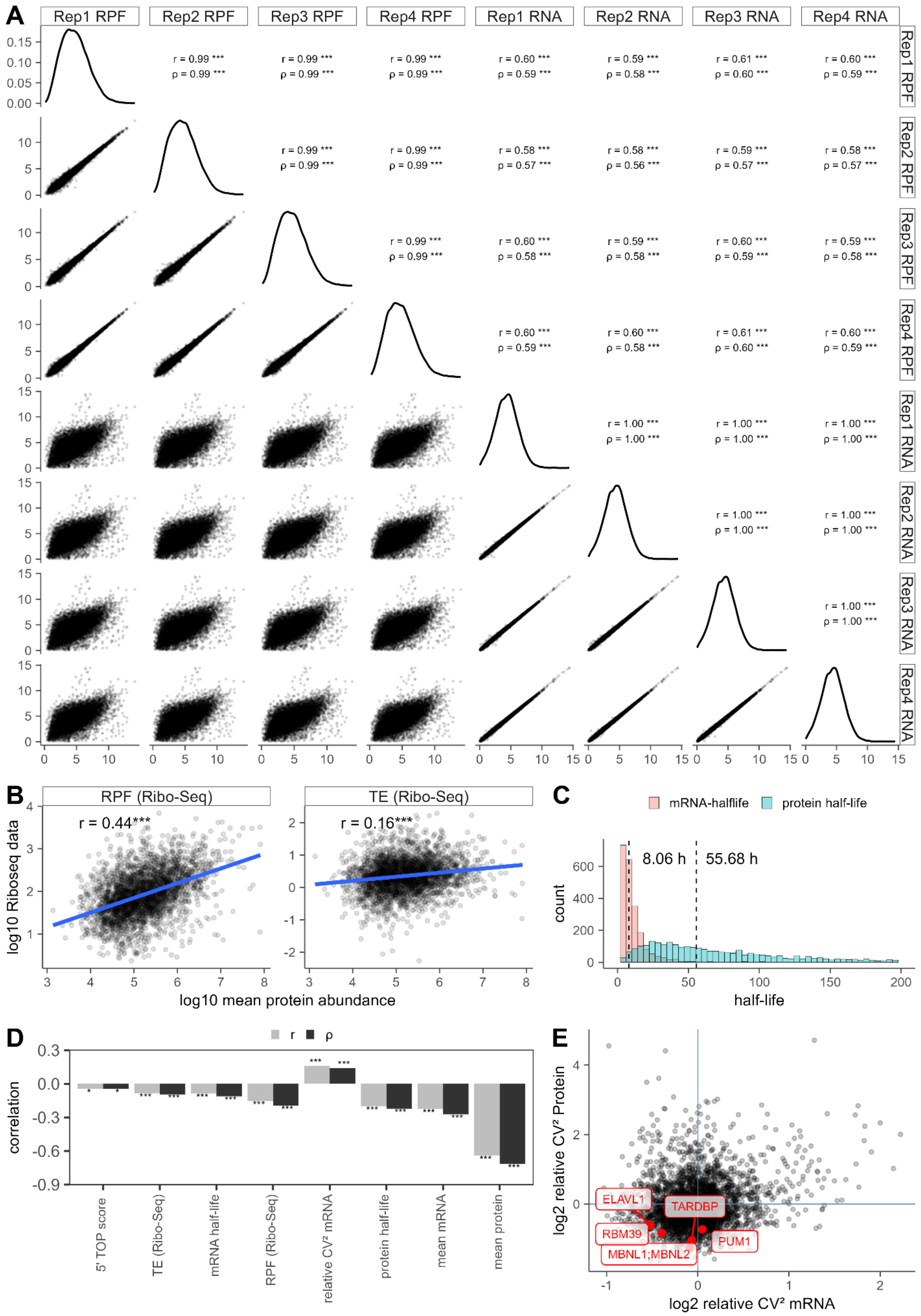


**Extended Data Fig. 5 | Ribosome profiling (Ribo-seq) and dynamic SILAC to assess proteome dynamics in bulk U2OS cells**

A-B | Ribo-Seq data

A) Correlation of Ribo-Seq replicates (log 2 tpms for RPFs and mRNA) used to calculate average RPF tpms, average mRNA tpms and Translation efficiency (TE). Pearson (r) and Spearman (rho) correlations are indicated in the upper quadrants. Density distribution of tpms is plotted in the diagonal.

B) Log 10 scaled RPF average tpms (left) and average Translation Efficiency (right) vs log10 mean protein abundance. Pearson correlation is indicated in the the plot, *** indicate a p-value < 0.0001.

C | Distribution of mRNA half-lifes and protein half-lifes.

mRNA half-lifes in U2OS cells are from Berg et al. 2024. Protein turnover in U2OS has been acquired on a Thermo Astral instrument in DIA mode and half-lifes have been calculated based on a cell cycle time derived from histone degradation rates (Leduc et al. 2025). The median mRNA half-life is at 8.06 h while the median protein half-life is at 55.68 h and thereby in similar ranges as in Schwanhäusser *et al*. (2011).

D | Correlations of $CV^2$ with different features.

Pearson and Spearman correlation of log2 $CV^2$ protein with a variety of different log2 transformed features, including mean protein abundance which is the main driver of not-normalized $CV^2$ values.

E | Relative protein and mRNA $CV^2$ and a couple of selected autoregulatory genes.

Log2 relative $CV^2$ for mRNA and protein (Extended Data Figure 3 A and Fig 3B). Blue lines separate the different noise category areas. Selected proteins are highlighted in red.

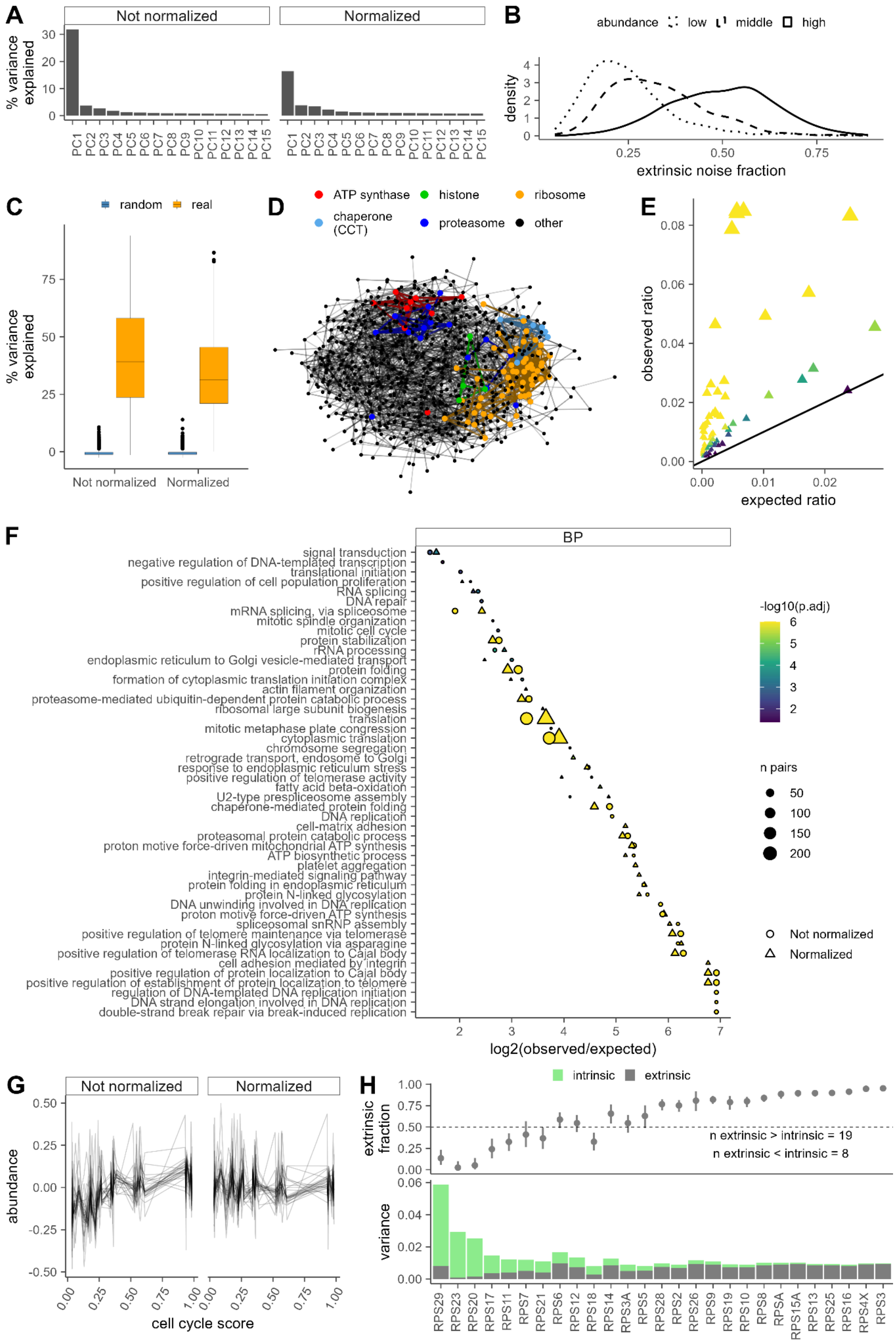


**Extended Data Fig. 6 | Protein noise is mainly extrinsic.**

A | Variance explained by PCA.
% of variance explained by the first 15 components of a PCA for not normalized (left) and cell cycle and cell size normalized (right) data. The first nine components for not normalized and the first 13 components for normalized data have been determined by parallel analysis to be significant. They have been assumed as indicator of cellular states and used to calculate the extrinsic noise fractions (Fig. 4A and Extended Data Fig. 6B).

B | Extrinsic noise fraction of cell cycle and cell size normalized data.
Extrinsic noise fraction based on the variance explained for a given protein by the first 13 components of the PCA given in A) for different protein abundance categories (lower third, dotted; middle third, dashed; higher third, continuous).

C | Comparing Ridge regression model quality to random models.
The network analysis in D) and E) as well as Figure 4B-D is based on Elastic Net regression predicting the abundance of a given protein by all other proteins for either normalized or not normalized data. As control, random models (blue) were trained.

D | Network for cell cycle and cell size normalized data.
Ridge regression based network of all proteins where the abundance could be modelled by the abundance of all other proteins with an $R^2$ of > 0.40 and that were among the top 15 predictors (only positive coefficients) of one of these proteins. Notes without edges have been removed. Members (nodes and shared edges) of selected protein complexes are highlighted.

E-F | GO enrichment.
For each protein in the network, the GO terms were pulled and the number of shared GO terms between a protein pair versus the total number of unique protein pairs (observed ratio) has been calculated. To verify the significance, a hypergeometric test was performed where the theoretical paired ratio of a given GO term within the whole network was calculated (expected ratio). The number of pairs per shared GO term is indicated by the dot size (minimum 4 pairs), the adjusted p-value (any p value lower than 10^-6 was set to that value and any value bigger than 0.06 was set to 0.06) acquired with the hypergeometric test is indicated by the colour of the dots. **E)** Observed ratio versus expected ratio of cell cycle and cell size normalized data for all GO terms (all ontologies) in the network. Line indicates the diagonal where observed = theoretical ratio. **F)** Significantly enriched biological process terms for not normalized (round) and normalized (triangle; Fig. 4C-D) data, x axis indicates log2 FC between observed and expected ratio.

G-H | Small Ribosomal Subunit.
Noise of proteins corresponding to the small ribosomal subunit. **G)** Median centered individual protein abundance traces over the cell cycle score for not normalized (left panel) and normalized (right panel) data. For better visibility, a randomly sampled subset of cells was used. **H)** Decomposition of extrinsic and intrinsic noise for non-normalized ribosomal proteins of the small subunit (RPSs). The upper panel shows the estimated fraction of total noise that is extrinsic, with confidence intervals estimated by bootstrapping. The lower panel displays the total variance, decomposed in extrinsic (grey) and intrinsic (green) components. Extrinsic variance is defined as the variance in the abundance of each RPL that can be

explained by the abundance of all other RPSs. Intrinsic variance corresponds to the residual variance that cannot be explained by the abundance of other RPSs.

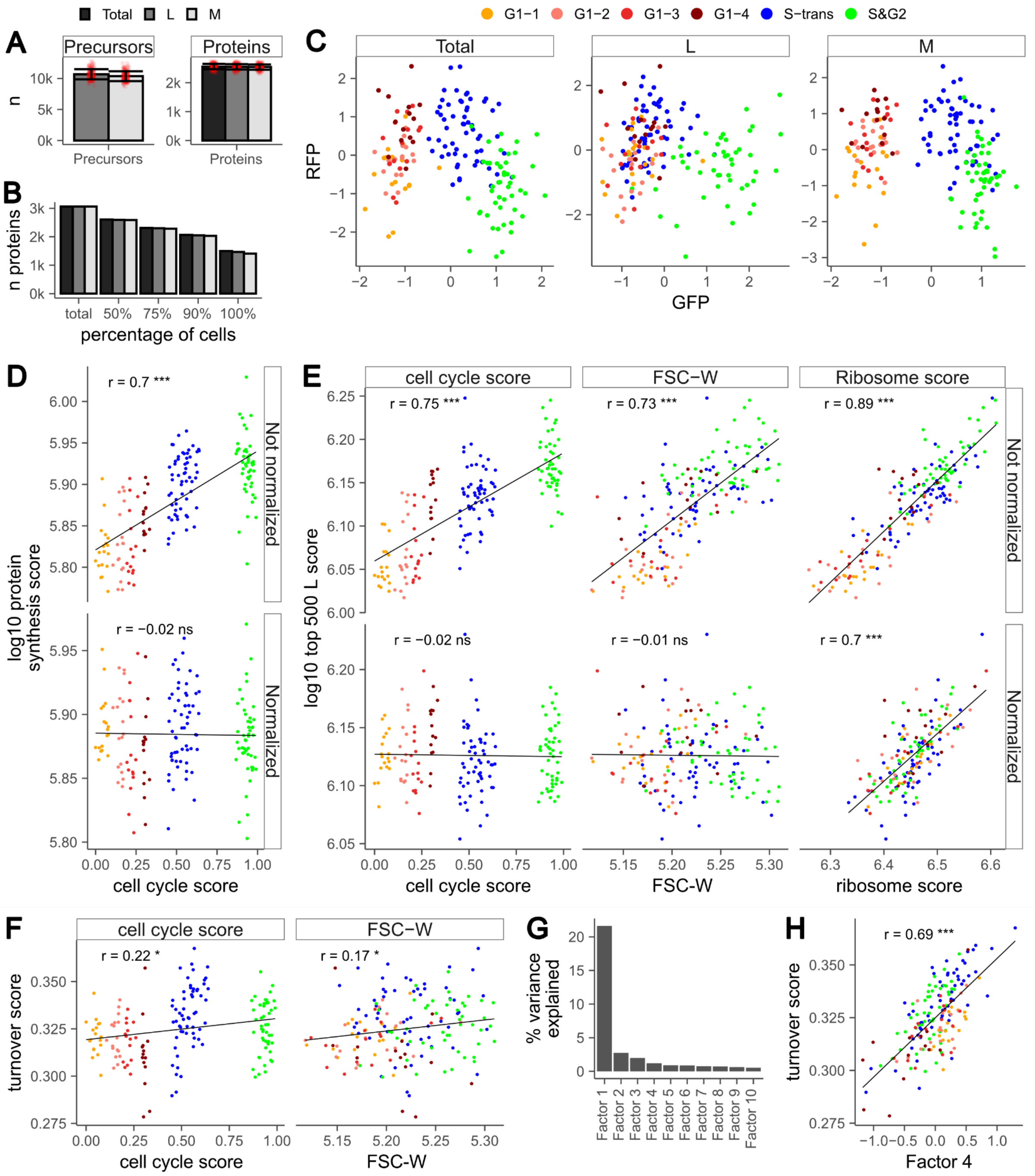


**Extended Data Fig. 7 | Quality and further analysis of pulsed single cell data set.**

A-B | Number of identifications and data completeness across channels**.**

Mean number of precursors and proteins (A) and their data completeness (B) across all 170 pulse SILAC single cell measurements. L: Old proteome, M: Newly synthesized proteome (within the last 9.5 h), Total: L+M. Error bars indicate standard deviation, red dots are individual runs. Y axis in thousands.

C | RFP and GFP behaviour across the cell cycle for the different channels.

Z-scored GMMN-GFP and CDT1-RFP intensities measured with mass spectrometry for the different proteomes. Notably, the newly synthesized proteins (M) are quantifying the accumulation of GMMN-GFP beginning in S-phase leading to a separation between S and

G1 phase cells (compare L to M). The total proteome shows the combined behaviour of L and M.

D | Protein synthesis score versus cell cycle

Log10 protein synthesis score as a proxy for translation over cell cycle score, for non-normalized (upper panel) and normalized (lower panel) data. The protein synthesis score is the median M SILAC abundance (the newly synthesized proteome) of the top 500 most abundant proteins that were quantified in all cells excluding ribosomal proteins as well as the subset of proteins that was used to calculate the random control. Significance is indicated as follows: * $p \leq 0.05$; ** $p < 0.001$; *** $p < 0.0001$.

E | Using top 500 L proteins instead of the protein synthesis score (top 500 M proteins)

As an alternative control to the randomly sampled proteins, the same top 500 proteins used to calculate the proteins synthesis score were used to calculate a score using only L SILAC abundances. These correspond to the pre-existing proteome. This control is shown for the non-normalized (upper panels) and the normalized data (lower panels) over the cell cycle score (left panels), log10 FSC-W as cell size (middle panels) and ribosome score (left panels). Significance is indicated as follows: * $p \leq 0.05$; ** $p < 0.001$; *** $p < 0.0001$.

F | Turnover score versus cell cycle and cell size.

The turnover score ( $\Sigma M / \Sigma(L+M)$ ), which resembles the fraction of medium-heavy intensities in relation to the total intensities per cell, is shown over cell cycle score (left panel) and cell size (right panel) for the non-normalized data.

G | Variance explained by MOFA factors

% variance explained by the first 10 MOFA factors.

H | Correlation of Factor 4 with turnover.

The turnover score over the low dimensional representation of the MOFA (Z) for Factor 4. Since signs in MOFA are arbitrary, the Factor 4 Z values have been multiplied with -1, leading to a positive correlation.
